# Rhizobial infection-specific accumulation of phosphatidylinositol 4,5-bisphosphate inhibits the excessive infection of rhizobia in *Lotus japonicus*

**DOI:** 10.1101/2025.06.26.661640

**Authors:** Akira Akamatsu, Toshiki Ishikawa, Hiroto Tanaka, Yoji Kawano, Makoto Hayashi, Naoya Takeda

## Abstract

- During the symbiosis of legume with nitrogen-fixing bacteria, collectively called rhizobia, suppression of excessive rhizobial infection by host plants is important to maximize the benefits of symbiotic nitrogen fixation. However, it remains relatively poorly understood the molecular mechanism involved in the suppression.
- We performed LC-MS and RNA-seq analysis using rhizobia-infected *Lotus japonicus* roots and investigated the role of phosphatidylinositol (PI) and phosphatidylinositol phosphates (PIPs) in the symbiosis. *Phosphatidylinositol transfer protein* (*PITP*)*-like proteins 4* (*PLP4*)*, phosphatidylinositol 3-phosphate 5-kinase 4* (*PIP5K4*) and *PIP5K6* mutants, which are involved in the vesicular transport of lipids and phosphorylation of PIPs, were used to show the involvement of the signaling of PI and PIPs. Accumulation of phosphatidylinositol 4,5-bisphosphate [PI(4,5)P_2_] during rhizobial infection were examined by a fluorescent marker 1xTUBBY-C (TUBBY).
- We found that PI signaling-related genes were upregulated, and the amount of PIP_2_ increased in *L. japonicus* roots during rhizobial infection. In the *PLP4*, *PIP5K4* and *PIP5K6* mutants, rhizobial infection increased and PIP_2_ accumulation was failed. Furthermore, the observation of PI(4,5)P_2_ in rhizobia-infected roots revealed that the ectopic accumulation was closely related to suppression of rhizobial infection.
- Our findings indicate that the accumulation of PI(4,5)P_2_, which is mediated by PLP and PIP5Ks, suppresses excessive rhizobial infection in the root epidermis and cortex, leading to the optimal number of nodules.

## Introduction

Nitrogen-fixing symbiotic bacteria, collectively called rhizobia, invade legume plant cells, resulting in formation of root nodules in which atmospheric nitrogen is converted to ammonia. The ammonia produced by rhizobia is an essential nutrient for host plant growth when soil nitrogen is poor. For this reason, root nodule symbiosis (RNS) has long been considered to be an agriculturally important trait.

During rhizobial infection, nodulation factors (Nod Factors: NFs), lipochito– oligosaccharides produced by rhizobia are important signals that trigger host plant symbiotic responses (Oldroyd, 2013). NF recognition by plasma membrane (PM) receptors in host plants induces rapid changes in Ca^2+^ concentration in and around the nucleus, termed Ca^2+^ spiking. Ca^2+^ spiking is decoded by Ca^2+^/calmodulin-dependent protein kinase (CCaMK), and CYCLOPS phosphorylated by CCaMK induces expression of two transcription factors, *Nodule Inception* (*NIN*) and ERF REQUIRED FOR NODULATION1 (ERN1) (Schauser *et al*., 1999; Singh *et al*., 2014; Cerri *et al*., 2017). In the model legumes *Lotus japonicus* and *Medicago truncatula,* rhizobia enter the host cell at the tip of root hairs. On receiving NF signals, the root hair tip curls to create an infection chamber, a space to enclose the rhizobia (Fournier *et al*., 2015). Rhizobia grow in the chamber, the stage of infection called a microcolony (Gage, 2004), and the cell membrane surrounding the rhizobia swells, creating polarity at the swelling site and initiating cell invasion (Liu *et al*., 2019). At the stage of infection thread formation in root hairs, cell division in the cortex is initiated, leading to the development of these cells into nodules (Suzaki et al., 2012). Rhizobia invade deeper into the root through tunnel-like structures called infection threads. Infection threads that progress from the epidermal to the cortical cells often branch their tips and eventually release rhizobia into the nodule cells (Rae *et al*., 2021).

Although RNS promotes host plant growth, excessive rhizobial infection significantly inhibits plant growth. In supernodulation mutants, numerous nodules are formed, whereas overall plant growth is impaired. This highlights the critical importance of maintaining a symbiotic balance between the host plant and rhizobial bacteria (Buttery, 1989; Wopereis *et al*., 2000; Ferguson *et al*., 2014). Rhizobial infection induces suppression of further infection to maintain an optimal level of symbiosis. The suppression system involves long-distance regulation, which regulates rhizobial infection via the shoot. The transcription factor NIN, which promotes RNS, also plays a role in the suppression system. In *L. japonicus* roots, CLE-RS1/2 peptides induced by NIN are transported from root to shoot via the xylem and are recognized by the HAR1 receptor (Okamoto *et al*., 2009; Okamoto *et al*., 2013; Soyano *et al*., 2014). In the absence of rhizobial infection, shoot-derived microRNA2111 suppresses the expression of the F-box protein TOO MUCH LOVE (TML) in roots. In contrast, during rhizobial infection, the expression of microRNA2111 is downregulated, resulting in increased TML expression and subsequent suppression of further rhizobial infection (Magori *et al*., 2009; Tsikou et al., 2018). This cascade is called autoregulation of nodulation (AON). The suppression of rhizobial infection is also regulated by plant hormones. Treatment of either ET precursor ACC or GA inhibit nodule formation (Oldroyd et al., 2001; Maekawa et al., 2009). *M. truncatula ethylene insensitive 2* (*Mtein2*, also referred to as *SICKLE*), in which the ET signaling is compromised, shows increase of the number of nodules (Penmetsa and Cook, 1997). In *L. japonicus* LjEIN2-1 and LjEIN2-2 cooperatively regulate the nodule number (Miyata *et al*., 2013). In addition, *M. truncatula* ent-copalyl diphosphate synthase 1 (CPS1) and GA3 oxidase 1 (GA3ox1), which are involved in the biosynthesis of bioactive GA, are transcriptionally regulated by NIN (Gao et al., 2023). In a cytokinin perception mutant *HYPERINFECTED 1* (*hit1*) the number of infection threads also increases, suggesting that cytokinin functions to suppress rhizobial infection (Murray et al., 2007). Thus, leguminous plants employ multiple mechanisms to restrict nodulation.

Phosphatidylinositol (PI), a membrane phospholipid with anionic polarity, undergoes phosphorylation on the inositol ring to become phosphatidylinositol phosphates (PIP and PIP_2_), such as PI(3)P, PI(4)P, PI(3,5)P_2_, and PI(4,5)P_2_, each with specific functions (Noack & Jaillais, 2020). The PIP and PIP_2_ are produced by the catalysis of specific kinases, including phosphatidylinositol 4-kinase (PI4K) and phosphatidylinositol-4-phosphate 5-kinases (PIP5K). In plants, phosphatidylinositol phosphates are known to be involved in the apical cell expansion of tip-growing cell, such as root hairs and pollen tubes (Ischebeck *et al*., 2008; Hirano *et al*., 2018). *Arabidopsis* PIP5K3 (AtPIP5K3) and its product PI(4,5)P_2_ localize to the apex of the root hair, and they are required for polar tip growth (Hirano *et al*., 2018). In *Arabidopsis*, the PI(4,5)P_2_ produced by AtPIP5K4 and AtPIP5K5 regulates the polar tip growth of pollen tubes via regulation of apical pectin deposition (Ischebeck *et al*., 2008). Furthermore, phosphatidylinositol phosphates are known to be involved in the host plant infection process by pathogenic microbes. In *Arabidopsis*, PI(4,5)P_2_ accumulates in the extrahaustorial membrane (EHM) of powdery mildew infection sites (Qin *et al*., 2020). Loss of function of PIP5Ks results in suppression of infection, indicating that PI(4,5)P_2_ accumulation is an essential factor in the infection of powdery mildew (Qin *et al*., 2020). It has also been shown that PI(4,5)P_2_ accumulation in the extra-invasive hyphal membrane (EIHM) is conducive to *Colletotrichum tropicale* infection, which causes plant anthracnose (Shimada *et al*., 2019). In addition to *C. tropicale*, during infection of the oomycete *Hyaloperonospora arabidopsis*, a causal pathogen of downy mildew, PI(4)P is enriched in the EHM (Shimada *et al*., 2019). When arbuscular mycorrhizal fungi establish symbiosis with *M. truncatula*, PI(4,5)P_2_ accumulates in periarbuscular membrane at the invading hyphae and at the arbuscule trunks (Ivanov & Harrison, 2019). These findings indicate that phosphatidylinositol phosphates are involved in plant-microbe interactions, especially when tip-growing hyphae invade into host cells.

Only little information is available for the behavior of phosphatidylinositol phosphates in rhizobial infection. Phosphatidylinositol 3 kinase (PI3K) has been shown to be involved in the generation of reactive oxygen species when rhizobia invade root hair cells (Peleg-Grossman *et al*., 2007; Robert *et al*., 2018). *LjPLP4* (formerly *PLP-IV*) encodes a protein with Sec14 and Nodulin domains in which the expression of the mRNA encoding the C-terminus of the protein (LjNOD16) is induced in nodules (Kapranov et al., 1997; Kapranov et al., 2001). The Sec14 domain of LjPLP4 has a PI transfer activity (Kapranov *et al*., 2001). In addition, AtSFH1, the orthologous gene of LjPLP4 in *Arabidopsis*, promotes the accumulation of PI(4)P and PI(4,5)P₂ by facilitating the functions of PI4K and PIP5K to the PM. However, it is still unknown whether phosphatidylinositol phosphates, especially PI(4,5)P_2_, are involved in rhizobial infection. Therefore, investigating phosphatidylinositol phosphate dynamics is helpful to understand how they play roles in RNS. In this study, we showed that increased expression levels of PI signaling-related genes led to accumulation of PI and phosphatidylinositol phosphates in a rhizobial inoculation-dependent manner. In the mutants of *LjPLP4*, *LjPIP5K4* and *LjPIP5K6,* accumulation of PI and phosphatidylinositol phosphates was not induced by rhizobial inoculation, resulted in promotion of rhizobial infection. PI(4,5)P_2_ marker analysis revealed that accumulation of PI(4,5)P_2_ correlated with suppression of rhizobial infection, and successful infection eliminated accumulation of PI(4,5)P_2_. The role of PI(4,5)P₂ in RNS appears to differ from its function in other plant–microbe interactions, potentially contributing to the fine-tuning of RNS through the suppression of excessive rhizobial infection.

## Materials and Methods

### Plant materials and growth conditions

We used the wild type *L. japonicus* MG-20, and symbiosis mutants in the MG-20 background (*har1-7*, *nin-9*, *cyclops-6,* and *tml*) which was described previously (Magori *et al*., 2009; Suzaki *et al*., 2012). The mutants produced by CRISPR/Cas9 in this study were *Ljplp4-4-10*, *Ljplp4-4-11*, *Ljplp4-9*, *Ljpip5k4-2*, *Ljpip5k4-3*, *Ljpip5k4-9*, *Ljpip5k6-1*, *Ljpip5k6-5*, and *Ljpip5k6-6* as described below. The *tml har1*, *tml Ljplp4-4-10*, *har1-7 Ljplp4-4-10* and *Ljplp4 YC2.60* mutants were produced by crossing. From the F_2_ population, the homozygous double mutants were selected using PCR. To produce the LjPLP4 OE, the full-length of *LjPLP4* driven by the Lotus Ubiquitin promoter (*pLjUbiquitin::LjPLP4*) were expressed in *L. japonicus* MG-20. The seeds were scarified with sandpaper, sterilized with sodium hypochlorite (effective chloride; approximately 1%) for 10 min, and soaked overnight in sterilized water. The sterilized seeds were germinated on 0.8% agar plates and grown in a growth chamber (24°C, 16-h light/8-h dark).

### Analysis of rhizobial infection and root hair growth

We used *Mesorhizobium loti* (MAFF303099), or transgenic *M. loti* strain carrying *DsRed* (Maekawa *et al*., 2009) and *GFP* (Akamatsu et al., 2022), for the quantitative analysis of infection threads and nodules. These strains were used to inoculate 3-day-old plants (3 days after germination, 2 days dark/1 day light) in plastic pots (SPL Life Sciences, Korea) containing 300 mL vermiculite supplied with B&D medium (Broughton, 1971) containing 0.1 mM KNO_3_ (100 mL per pot). A total volume of 1 mL *M. loti* solution with an absorbance of 0.1 at 600 nm was added to each pot. Root nodule infection phenotypes, including the number of nodules, infection threads, and infection events, were determined by roots inoculated with *M. loti* carrying DsRed. The number observed for each plant was shown. In the measurement of infection threads, the total number of microcolonies (MCs), infection threads (ITs), and ramifying ITs was shown (Figs. 2c, e, 4e, g, 5e, g, i).

**Figure 1.**
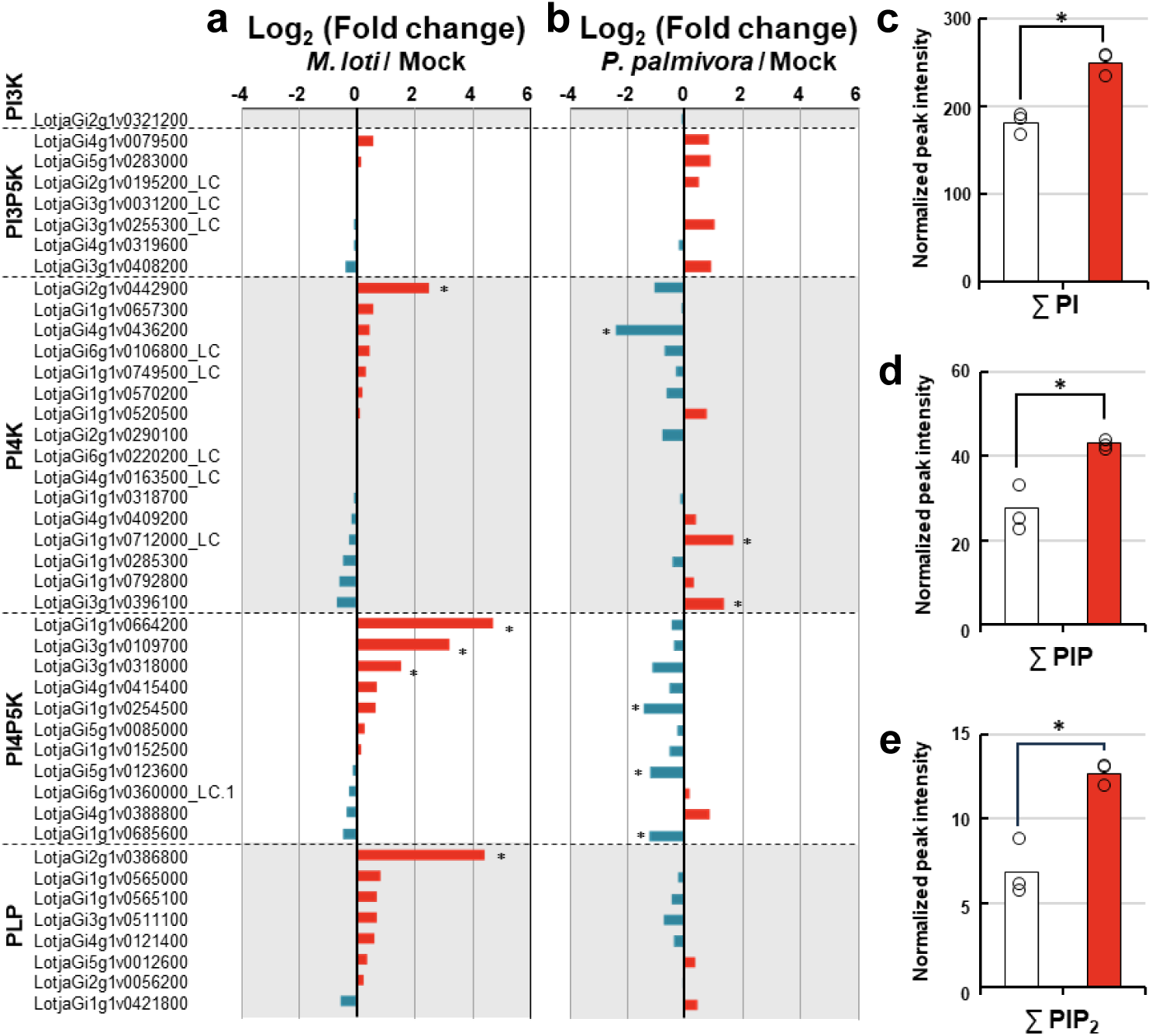
Expression of PI signaling-related genes and accumulation of PI, PIP, and PIP_2_. **a, b**. RNA-seq analysis using *Lotus japonicus* MG-20 roots during symbiotic or pathogenic microbe interaction. Relative expression in log_2_ (FC) of (*M. loti* / Mock) at 1 week after inoculation (wai) (a) or (*P. palmivora* / Mock) at 5 days after inoculation (dpi) (b). (*; log_2_(FC) ≥ 1 or log_2_ (FC) ≤ −1 and P ≤ 0.05). Red bars indicate increased expression, and deep blue bars indicate decreased expression. **c–e**. LC-MS analysis of phosphoinositol (PI) (c), phosphatidylinositol monophosphate (PIP) (d), and phosphatidylinositol bisphosphate (PIP_2_) (e) in *L. japonicus* roots under control conditions (white) or inoculated with *M. loti* for 1 week (Red) (n = 3 biological replicates, the bars show average, and circles show each data point, *t*-test; *P < 0.01).

**Figure 2.**
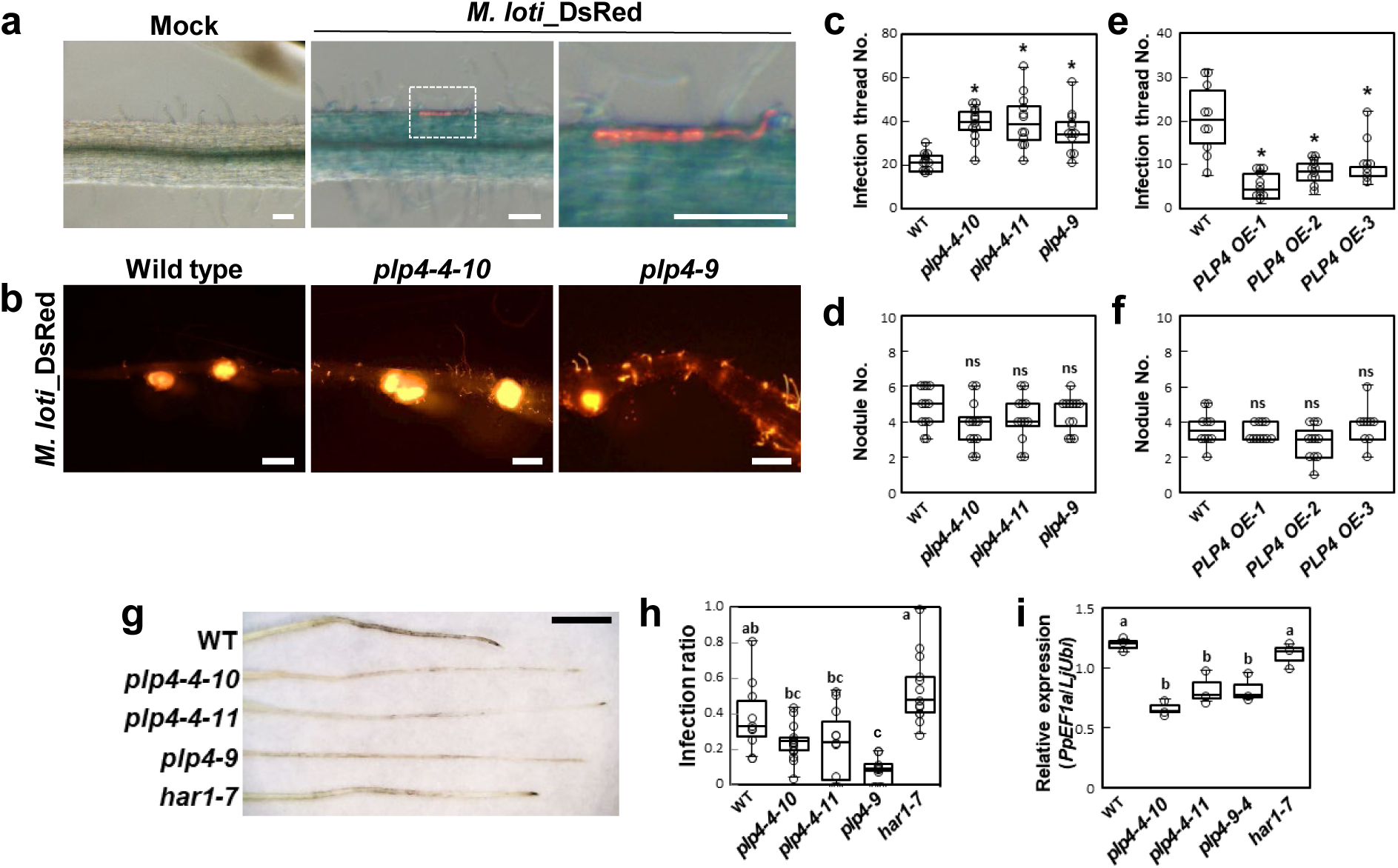
*LjPLP4* negatively regulates rhizobial infection but is required for *P. palmivora* infection. **a.** Rhizobia-infected and non-infected *L. japonicus* MG-20 wild type (WT) roots transformed with *pLjPLP4::GUS*. The roots were stained with GUS staining buffer at 2 wai of *M. loti* carrying DsRed. The image on the right is an enlargement of the framed area of the center image. Scale bars, 100 µm. **b–f.** The plants were inoculated with *M. loti* carrying *DsRed* for 1 week. **b**. Nodules and infection threads observed in wild type, *Ljplp4-4-10,* and *Ljplp4-9*. Scale bars, 200 µm. **c–f.** Quantitative analysis of rhizobial infection phenotypes of *Ljplp4* mutants (c, d) and *LjPLP4* overexpressing (OE) plants (e, f). n ≥ 12, Dunnett’s test, one asterisk (*) indicates *P* < 0.01. ns: not significant. Similar results were obtained in more than three independent experiments. **g–i.** *P. palmivora* infection phenotypes in *L. japonicus* MG-20 wild type, *Ljplp4-4-10*, *Ljplp4-4-11*, *Ljplp4-9,* and *har1-7* at 5 dpi. **g**. Representative image of *P. palmivora* infected *L. japonicus* roots in *LjPLP4* and *har1-7* mutants. Scale bar, 0.5 cm. **h, i.** Quantitative analysis of *P. palmivora* infection. Infection ratio (infected lesion/root length) (h). Measurement of *PpEF1a* expression in *P. palmivora* infected roots by real-time PCR (i). n ≥ 12, Tukey test, *P* < 0.01. Similar results were obtained in three independent experiments.

**Figure 3.**
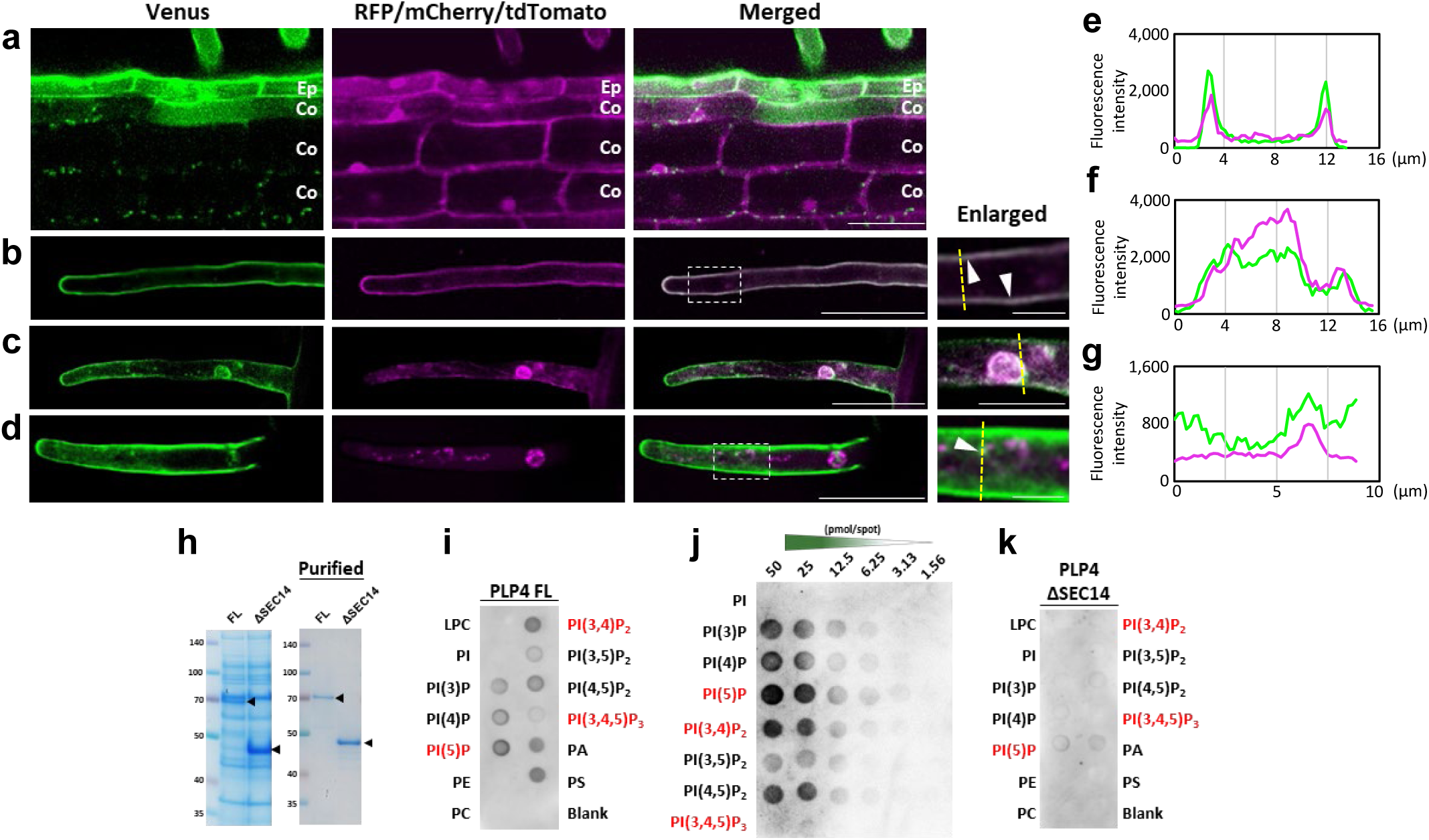
The plasma membrane localization of LjPLP4 in *L. japonicus* roots. **a–d.** Hairy roots transformed with the fluorescent constructs. **a**. Venus-LjPLP4 (green) and Free RFP (magenta), which is a transformation marker, were expressed in *L. japonicus* wild type roots. The images show the epidermis and the cortex. Abbreviations are Ep, epidermis; Co, cortex. **b–d**. Venus-LjPLP4 (green) was co-expressed in root hair cells with NFR1-mCherry (magenta) (b), tdTomato-CTT (magenta) (c), and GmMAN49-mCherry (magenta) (d). Scale bars, 50 µm or 10 µm in the enlarged images. The arrowhead indicates a merged signal of green and magenta. **e-g**. Intensity profiles of Venus and RFP/mCherry/tdTomato signals along the yellow transect shown in (b-d). **h**. CBB images of purified HIS-LjPLP4 full length (FL) or ΔSEC14. **i-k**. PIP-strip, membrane binding assays with lipid-spotted membranes. **j**. PIP Arrays, membranes pre-spotted with a concentration gradient. The membranes were incubated with 1.0 µg/mL of LjPLP4 FL (i, j) or ΔSEC14 protein (k). The lipids shown in red are not present in plants. Abbreviations are LPC, lysophosphatidylcholine; PE, phosphatidylethanolamine; PS, phosphatidylserine; PC, phosphatidylcholine.

**Figure 4.**
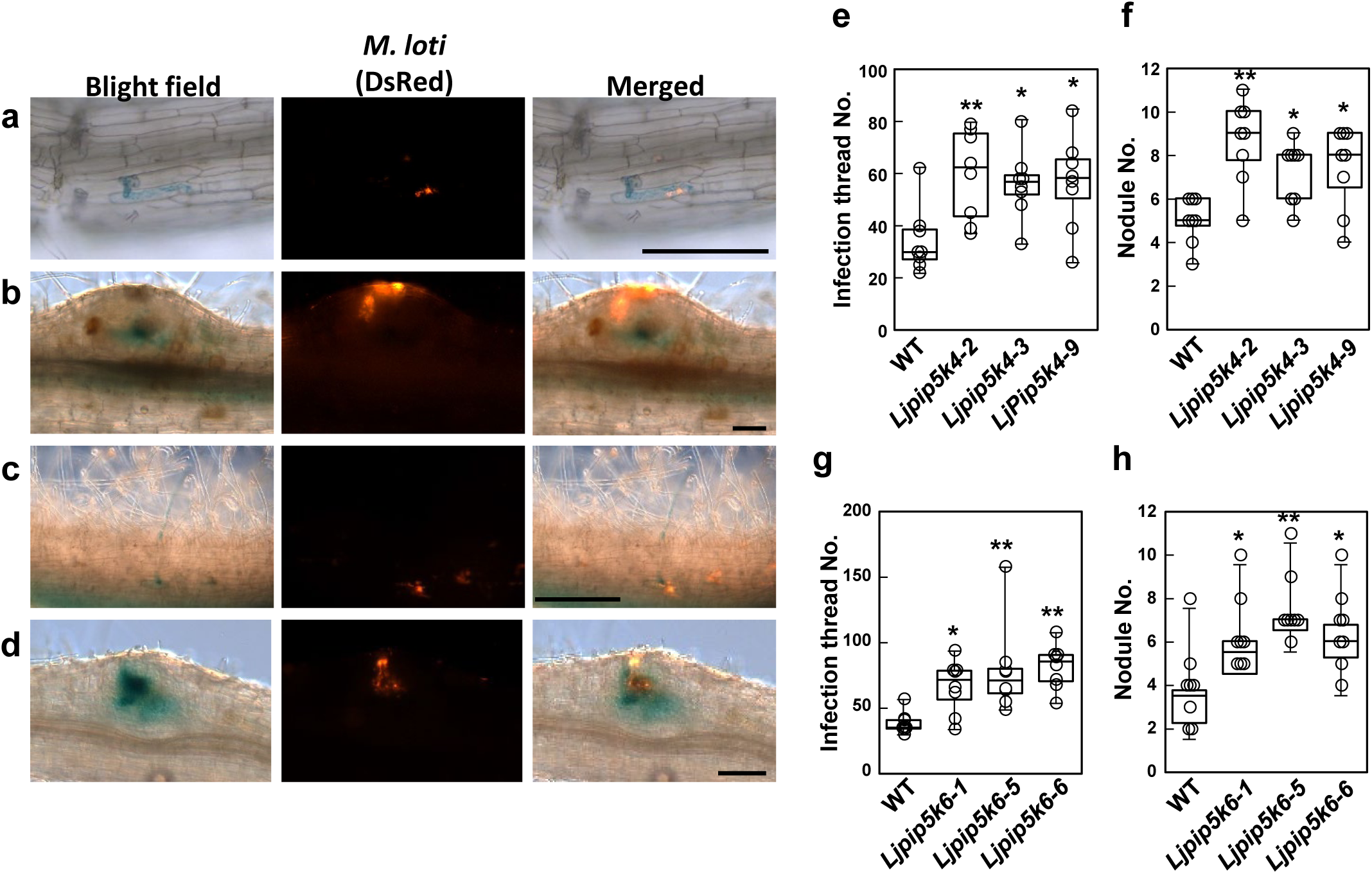
*LjPIP5K4* and *LjPIP5K6* negatively regulates rhizobial infection. **a.** Rhizobia-infected *L. japonicus* MG-20 wild type (WT) roots transformed with *pLjPIP5K4::GUS* (a, b) or *pLjPIP5K6*::*GUS* (c, d). Infected epidermal cells (a, c), nodule primordium (b, d). The roots were stained with GUS staining buffer at 2 wai of *M. loti* carrying DsRed. Scale bars; 100 µm. **e-h.** Quantitative analysis of rhizobial infection phenotypes of *Ljpip5k4* and *Ljpip5k6* mutants. The plants were inoculated with *M. loti* carrying *DsRed* for 1 week (n ≥ 10). Dunnett’s test, *P < 0.05, **P < 0.01, ns; not significant. Similar results were obtained in more than three independent experiments.

**Figure 5.**
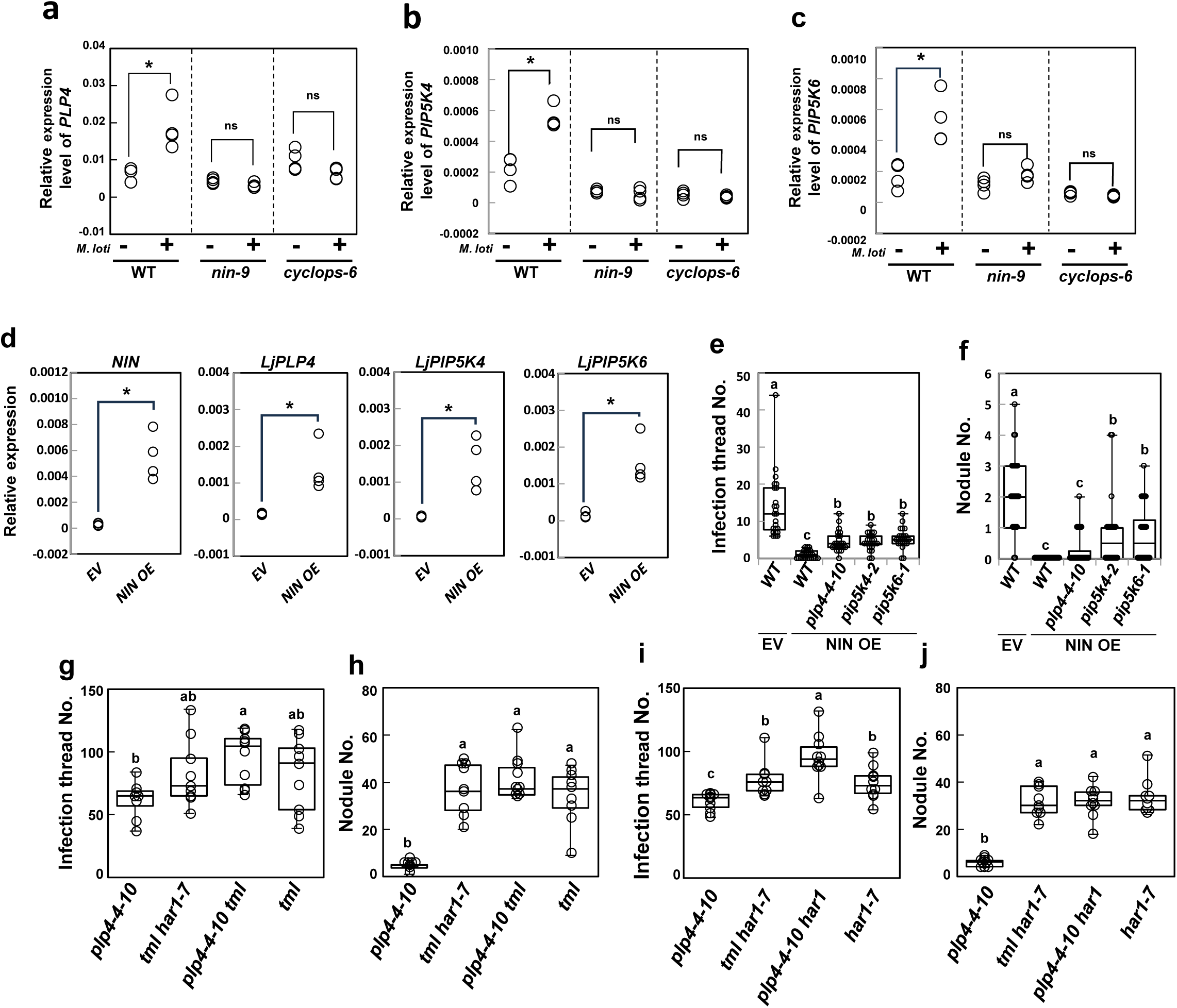
*NIN* regulates the suppression of rhizobial infection by *LjPLP4*, *LjPIP5K4* and *LjPIP5K6*. The plants were inoculated with *M. loti* wild-type (a-c) or carrying DsRed (e-i) for 1 week. **a-c**. Measurement of *LjPLP4*, *LjPIP5K4* and *LjPIP5K6* expression in *nin-9* and *cyclops-6* mutants by real-time PCR (n = 4, t-test, *P < 0.05, ns; not significant). **d**. Measurement of *NIN, LjPLP4, LjPIP5K4* and *LjPIP5K6* expression in *NIN*-overexpressing roots by real-time PCR (n = 4, t-test, *P < 0.01). **e-f**. Quantitative analysis of rhizobial infection phenotypes of *NIN*-overexpressing roots in wild type, *Ljplp4*, *Ljpip5k4* and *Ljpip5k6* mutants, and wild type roots transformed with empty vector (EV). **g-j**, *plp4-4-10*, *tml har1-7*, *plp4-4-10 tml* and *tml* (g, h), or *plp4-4-10*, *tml har1-7*, *plp4-4-10 har1-7* and *har1-7* (i, j). n ≥ 12, Tukey test, *P* < 0.05. Similar results were obtained in more than three independent experiments.

To measure the root hair length in *L. japonicus*, root hairs located 5 mm from the root tip of seedlings (2 days after germination) were imaged under a upright microscope (Leica DFC7000 T camera mounted on a Leica DM6 B). The images obtained were used directly for root hair length measurement using ImageJ software (https://imagej.net/ij/).

### *Phytophthora palmivora* infection and phenotypic analysis

*P. palmivora* (E.J. Butler) (Rahman *et al*., 2014) was grown on V8 juice Original (Campbell’s, USA) agar medium for 6–10 days until mycelium was fully expanded in the 90-mm Petri dishes. The plates were kept in a fume hood for 24 h to dry the medium. A total volume of 10 mL sterilized cool water was poured on each Petri dish, allowed to sit for 1 h to induce the release of zoospores, and the concentration of zoospores was quantified on a hemacytometer. *L. japonicus* MG-20 wild-type and mutant seedlings were grown on 0.8% agar medium (3 days after germination, 2 days dark/1 day light), then inoculated at 23℃ with or without 1 mL (1 × 10^6^ zoospore/mL) of *P. palmivora* solution. To quantify *P. palmivora* growth on the roots, roots 5 days post-inoculation were used to measure the lesion size, which was normalized to individual root length. These same root samples were then frozen in liquid nitrogen to extract RNA for real-time PCR analysis using the *P. palmivora* EF1a gene compared with the *L. japonicus Ubiquitin* housekeeping gene.

### Plasmid construction

Gene fragments used to create binary vectors for plant transformation and protein expression in *Escherichia coli* were amplified by PCR using Prime STAR GXL DNA polymerase (Takara, Japan), and the specific primer sets were listed in Table S1. The promoter region (3 kbp) of *LjPLP4*, *LjPIP5K4* or *LjPIP5K6* was amplified using the primer set 1, 2 or 3 (Table S1) from the *L. japonicus* MG-20 wild-type genome, followed by cloning into the entry vector pENTR D/TOPO (ThermoFisher, USA). For the promoter-GUS fusion construction, the entry clones were transferred into the destination vector pKGWFS7 (Karimi *et al*., 2002) by the LR reaction. To construct the *pLjUbiquitin::Venus-LjPLP4*, *pLjUbiquitin::mCherry-LjPLP4*, *pLjUbiquitin::LjPIP5K6-Venus*, or *pLjUbiquitin::NFR1-mCherry* binary vectors, the coding sequence (CDS) of *LjPLP4*, *LjPIP5K6* or *NFR1* was amplified from cDNA synthesized from total RNA of *L. japonicus* MG-20 wild type using specific primer sets 4, 5, 6, or 7 (Table S1). They were fused with the amplified entry vector containing Venus or mCherry by inverse PCR using the primer set 8, 9, 10, and 11 (Table S1), using the SLiCE method (Motohashi, 2017). For the overexpression construction, the entry clones were transferred into the destination vector *pLjUbiquitin* (Maekawa *et al*., 2008) by the LR reaction. To express HIS-tagged LjPLP4 protein in *Escherichia coli*, *LjPLP4* was amplified using the specific primer set 12 and cloned into amplified pET28b vector using the PCR using the primer set 13 combined with the SLiCE method. To express HIS-tagged LjPLP4 ΔSEC14, pENTR-LjPLP4 ΔSEC14 was generated using the SLiCE method combined with Inverse PCR based on pENTR-LjPLP4. Based on this, LjPLP4 ΔSEC14 was amplified using the primer set 14 and fused with the pET28b vector using the SLiCE method.

### Real-time PCR analysis

Total RNA was extracted from root samples from 10 plants using the RNeasy Plant Mini Kit (QIAGEN, Netherlands). For all analyses, four or more independent biological replicates were obtained through separate samplings. The RNA concentration was quantified using NANODROP ONE (ThermoFisher, USA). Reverse-transcription and real-time PCR were performed using the ReverTra Ace qPCR RT kit (Toyobo, Japan) and the Thunderbird qPCR Mix (Toyobo, Japan) with the AriaMx Real-Time PCR System (Agilent, USA) according to the manufacturer’s instructions. The cDNA was synthesized from 500 ng total RNA, and each reaction was performed in triplicate to correct technical errors. The primer sets used to amplify *LjUbi*, *PpEF1a*, *CLE-RS1, LjPLP4, PIP5K4* and *PIP5K6* were listed in Table S1 (primer sets 15-20). Relative transcript levels were compared to *Ubiquitin* expression using the Ct value. Three to four biologically independent experiments were performed to calculate averages, and the statistical analysis compared with the control expression levels was performed using either Student’s t-test or Dunnett’s test.

### RNA-Seq analysis

*L. japonicus* MG-20 wild type was inoculated with *M. loti* (for 1 week) or *P. palmivora* (for 5 days) suspended in water on 0.8% agar. Also, for the negative control, roots were treated with water, and this treatment was designated as the mock condition. Total RNA was extracted from root samples from 10 plants using the RNeasy Plant Mini Kit (QIAGEN, Netherlands). To compare expression levels, total RNA was also extracted from plants that were not inoculated with *M. loti* or *P. palmivora*. Three biological replicates were conducted for each condition. RNA libraries from *L. japonicus* MG-20 roots that were infected were prepared for sequencing using the TruSeq Stranded Total RNA (Illumina, San Diego, CA, USA). Whole transcriptome sequencing was applied to RNA samples using the Illumina NovaSeq 6000 platform in the 101-bp single-end mode. Sequenced reads were mapped to the Gifu_1.2 genome (Kamal *et al*., 2020) using TopHat ver. 2.0.13 in combination with Bowtie2 ver. 2.2.3 and SAMtools ver. 1.0 (Li *et al*., 2009; Langdon, 2015). The number of fragments per kilobase of exon per million fragments mapped was calculated using Cufflinks ver. 2.2.1 (Trapnell *et al*., 2010).

### LC-MS analysis

Approximately 120 seedlings of *L. japonicus* MG-20 wild type, *Ljpip4*, and *Ljpip5k6*, were inoculated with *M. loti* suspended in water (for 1 week) on 0.8% agar plates, respectively. Also, for the negative control, roots were treated with water. PI, PIP, and PIP_2_ were analyzed by a phosphate methylation method coupled with LC-MS according to previous reports (Clark *et al*., 2011; Cai *et al*., 2016). In brief, lyophilized tissues (∼5 mg dry weight) were homogenized in 600 µl methyl-tert-butyl methyl ether (MTBE)/methanol (1:1, by vol). The homogenate was phase-partitioned by adding 700 µl MTBE and 300 µl 0.1 N HCl, and the upper ether layer was collected and dried using a centrifugal concentrator. The residue was dissolved in 200 µl of the upper phase of MTBE/methanol/0.01 N HCl (10:3:3, by vol) and mixed with 20 µl of (trimethylsilyl) diazomethane (2M in hexane, Sigma). Methylation of the phosphate groups was conducted for 20 min at room temperature and stopped by adding 6 µl acetic acid. After dryness, the residue was dissolved in methanol/acetonitrile/2-propanol (1:1:1 by vol) and analyzed by LCMS-8030 (Shimadzu) equipped with a Shim-pack Scepter C18 column (3 µm × 2.0 mm × 75 mm). The column was kept at 40°C and eluted with a binary gradient of solvent A (5 mM ammonium formate and 0.1% formate) and B (methanol/acetonitrile/2-propanol, 1:1:3 by vol, containing 5 mM ammonium formate and 0.1% formate): 0 to 15 min, 30%B to 100%B and maintained for 3 min. Permethylated PIPs were detected by the multiple reaction monitoring with [M+NH4]+ to [diacylglycerol+H−H2O]+. Lipid quantity was shown as normalized peak intensities using 18:0-20:4 PI (Sigma-Aldrich, USA) as an internal standard added to samples at the first extraction. Peak identity was confirmed using authentic standards of 18:1 PIP and 18:1 PIP_2_ (Sigma-Aldrich, USA).

### Hairy root transformation

The binary vectors were introduced into *L. japonicus* MG20 wild type or mutant plants by hairy root transformation using *Agrobacterium rhizogenes* AR1193 (Offringa *et al*., 1986). *A. rhizogenes* colonies were suspended in sterilized water. Roots of 6-day-old seedlings (3 days dark followed by 3 days in 16-h light/8-h dark, 24℃) were cut off below the hypocotyl in *A. rhizogenes* suspension. The infected shoots were incubated for 5 days in a growth chamber (24℃, 16-h light/8-h dark) on B5 medium (FUJIFILM, Japan) solidified with 1.0% agar and subsequently transferred to the same medium containing 12.5 μg/mL Meropen (Sumitomo Pharma, Japan) for 10 days to 3 weeks until bacterial growth was suppressed. Transgenic hairy roots were selected based on fluorescence under a stereomicroscope (SZX16; EVIDENT, Japan).

### Production of *Ljplp4, Ljpip5k4* and *Ljpip5k6* mutants using the CRISPR/Cas9 system, overexpression plants of *LjPLP4* and *UBQ10prom*::*2xCHERRY-1xTUBBY-C* (*TUBBY*) expressing plants

We used the CRISPR/Cas9 system to create mutation within the *LjPLP4* (LotjaGi2g1v0386800), *LjPIP5K4* (LotjaGi1g1v0664200) or *LjPIP5K6* (LotjaGi3g1v0109700) genes in the *L*. *japonicus* MG-20 genome. Two target sites were selected on the first exon and second exon of *LjPLP4* and the first exon for *LjPIP5K4* and *LjPIP5K6* using the CRISPR-P program (http://cbi.hzau.edu.cn/crispr/) (Lei *et al*., 2014) (Figs. S3a, S8a and S9a). The target oligonucleotides (primers set 21-22 for *LjPLP4*, 23-24 for *LjPIP5K4* or 25-26 for *LjPIP5K6* in Table S1) were annealed and cloned into the *Bbs*I site of the single guide RNA (sgRNA) vector pUC19_AtU6oligo (Ito *et al*., 2015). One sgRNA cassette was amplified using phosphorylated primers set 27 (Table S1), and the amplicon was ligated with the PCR product of the other sgRNA, including the pUC19 vector, amplified using the primer set 28 (Table S1), thereby constructing a vector in which two sgRNA cassettes were arranged in tandem. The sgRNA cassettes were subcloned into the I-SceI site of pZH_gYSA_FFCas9, which contained the Cas9 and HPT expression cassettes (Stiller *et al*., 1997; Ito *et al*., 2015; Nishida *et al*., 2018). The CRISPR/Cas9 constructs were introduced into *L*. *japonicus* MG-20 by *Agrobacterium tumefaciens* AGL1 as described previously (Stiller *et al*., 1997). The deletion or mutation of the target sites in the transgenic plants was examined by PCR using the primer set 29 for *LjPLP4*, 30 for *LjPIP5K4* or 31 for *LjPIP5K6* (Figs. S3b, S8b and S9b; Table S1) and confirmed by sequencing of the region (Figs. S3c, S8c and S9c). Homozygous lines of the deletion mutants were selected from the T1 transgenic plants, and the progeny (T2) were used in this study. For overexpression of *LjPLP4*, the *pLjUbiquitin::LjPLP4* constructs were introduced into *L*. *japonicus* MG-20 by *Agrobacterium tumefaciens* AGL1. The transgenic plants were examined by PCR using the primer set 32 (Table S1), and the expression level of *LjPLP4* was calculated by real-time PCR using the primer set 18 (Table S1). Homozygous overexpression lines were selected from the T2 transgenic plants and used in this study. TUBBY expressing plants were generated by transformation with AGL1 harboring AtUBQ10prom::2xCHERRY-1xTUBBY-C (TUBBY) (Simon *et al*., 2014) as described above.

### GUS staining

Transgenic hairy roots carrying the promoter GUS fusions were stained with a GUS staining buffer solution (0.5 mg/mL 5-bromo-4-chloro-3-indolyl-β-D-glucuronide [X-Gluc], 100 mM sodium phosphate buffer [pH 7.0], 100 mM EDTA, 0.5 mM K_4_[Fe(C.N.)_6_], 0.5 mM K_3_[Fe(C.N.)_6_], 0.1%v/v) at 30°C overnight. The stained roots were observed using a stereomicroscope (SZX16; EVIDENT, Japan).

### Subcellular localization and observation of PI(4,5)P₂

*Venus-LjPLP4* or *LjPIP5K6-Venus* expressing roots obtained using hairy root transformation were mounted on glass slides. As organelle markers, *pLjUbiquitin::NFR1-mCherry*, *p35S::tdTomato-CTT*, or *pMtBCP1::GmMAN49-mCherry* were co-expressed, respectively. Observations were performed under a confocal microscope A1 (Nikon, Japan) using a 20× objective lens. Observation of PI(4,5)P_2_ was performed using a confocal microscope SP8 (Leica Microsystems, Germany) using a 20× or 40× objective lens at 7 to 10 days after infection with *M. loti* carrying DsRed. All images in this study were processed with the software ImageJ.

### Grafting

The grafting experiment was performed as described by Magori et al. (2009). Three-day-old seedlings (MG-20 wild type and *Ljplp4*) were cut at the hypocotyls with a scalpel blade (Feather). Shoot scions were carefully cut at an angle and inserted into vertical slits made in the rootstock. The grafted plants were placed on pre-wet filter paper in a petri dish and covered with an additional filter paper. They were grown for 5 days under a 24°C condition with a 16-hour light/8-hour dark cycle. Then, plants were inoculated with *M. loti* carrying DsRed as described above, and infection was monitored one week later.

### Protein extraction and PIP strip

HIS-tagged fusion proteins were expressed in *E. coli* BL21 (DE3) and extracted using CelLytic ™ B Cell Lysis Reagent (Sigma-Aldrich, USA) added 8 M urea. Protein purification with HIS-Select® Spin Columns (Sigma-Aldrich, USA) was performed according to the manufacturer’s protocol. [Equilibration Buffer and Wash Buffer; 0.1 M sodium phosphate and 8 M urea (pH 8.0), Elution Buffer; 0.1 M sodium phosphate and 8 M urea (pH 4.5)]. PIP Strips (Echelon Biosciences, USA) were blocked with Blocking One (Nacalai Tesque, Japan) overnight at 4℃ in a Petri dish. The protein–lipid overlay incubation was performed with 1 mL HIS-tagged protein (1.0 µg/mL protein in PBS) at 4℃ overnight. The strip was washed three times for 10 min with TBS-T (50 mM Tris, 138 mM NaCl, 2.7 mM KCl, 0.1% Tween 20), then incubated with 1:2000 diluted His-tagged Polyclonal antibody (Proteintech, USA) in Solution 1 of Can Get Signal® Immunoreaction Enhancer Solution (TOYOBO, Japan) for 1 hour at room temperature. The strip was washed three times for 10 minutes with TBS-T and incubated for 1 h with 1:5000 Goat Anti-Rabbit IgG H&L (HRP) antibody (Abcam, UK) in Solution 2 of Can Get Signal® Immunoreaction Enhancer Solution. After repeating the wash step three times with TBS-T for 10 min each, protein–lipid binding was detected by ImageQuant 800 (GE Healthcare, USA).

### Statistical analysis

The data were analyzed using BellCurve for Excel (BellCurve, Japan). One-way ANOVA with Tukey test (Figs. 2h, i, 5e-h and S12a, b), two-way ANOVA with Tukey test (Fig. S13) or Dunnett’s test (Figs. 2c-f, 4e-h, S3d, e, S4, S5, S8d, e, S9d, e and S10) were applied. Student’s t-test by two-side were carried out (Figs. 1c-e, 5a-d, S1b and S12c, d). At least three biological repetitions were performed for all experiments.

## Results

### Expression of PI signaling-related genes and accumulation of PI and phosphatidylinositol phosphates are induced during rhizobial infection

To investigate PI and phosphatidylinositol phosphate dynamics during rhizobial infection (*M. loti*), an RNA-seq analysis was performed using *L. japonicus* roots 1 week after inoculation (Fig. 1a). We focused on the expression of PI signaling-related genes; *phosphoinositide 3-kinase* (*PI3K*) and *phosphoinositide 4-kinase* (*PI4K*), which phosphorylate phosphatidylinositol to phosphatidylinositol monophosphate (PIP), *phosphatidylinositol 3-phosphate 5-kinase* (*PI(3)P5K*) and *phosphatidylinositol 4-phosphate 5-kinase* (*PI(4)P5K* or *PIP5K*), which phosphorylate PIP to phosphatidylinositol bisphosphate (PIP_2_), and *phosphatidylinositol transfer protein* (*PITP*)*-like proteins* (*PLP*). The expression levels of *LjPI4K* (LotjaGi2g1v0442900) and three *PIP5Ks* (LotjaGi1g1v0664200, LotjaGi3g1v0109700, and LotjaGi3g1v0318000) were significantly upregulated by rhizobial infection (Fig. 1a). Due to specific PI(4,5)P_2_ or PI(4)P accumulation in *Arabidopsis* infected by powdery mildew or oomycete downy mildew, respectively, we performed another RNA-seq analysis during infection of *Phytophthora palmivora*, a pathogenic oomycete for *L. japonicus* (Rey *et al*., 2015; Fuechtbauer *et al*., 2018) (Fig. 1b). The expression of the PI signaling-related genes was generally suppressed upon infection with *P. palmivora*, especially that of *PIP5Ks*, which was markedly upregulated during rhizobial infection. Moreover, no PLP genes were significantly upregulated during *P. palmivora* infection. These results suggest that there is a difference in the role of PI signaling between rhizobial infection and pathogen infection of *P. palmivora*. To quantify the amounts of PI, PIP, and PIP_2_ in *L. japonicus* roots during rhizobial infection, we performed LC-MS. The results showed that accumulation of PI, PIP, and PIP_2_ was induced in a rhizobial infection-dependent manner (Fig. 1c-e). We initially focused on *LjPLP4* (LotjaGi2g1v0386800), which expression was only significantly induced among homologous genes for a PI transfer protein affecting the accumulation of PI, PIP and PIP_2_ (Fig. 1a). Quantitative polymerase chain reaction (PCR) confirmed that *LjPLP4* is the only *PLP* gene whose expression is upregulated during rhizobial infection among the eight *PLP* genes in the *L. japonicus* genome (Fig. S1).

### *LjPLP4* suppresses rhizobial symbiosis while it has a promoting role in oomycete infection

To confirm the tissue-specific expression of *LjPLP4* during rhizobial infection, we performed GUS analysis with the promoter region of *LjPLP4* and found that the GUS signal was induced by rhizobial infection throughout the root compared with roots that were not infected with rhizobia, but not in nodules (Figs. 2a and S2). We generated *LjPLP4* mutants using the CRISPR/Cas9 system to investigate whether *LjPLP4* was required for rhizobial infection (Fig. S3). None of these mutants had any effect on root growth (Fig. S3d-g). Inoculation of *LjPLP4* mutants with rhizobia resulted in a marked increase in the number of infection threads when compared with the wild type (Fig. 2b-c). Moreover, no significant difference was observed in the number of nodules compared with the wild type (Fig. 2d). We investigated at which stage of rhizobial invasion *LjPLP4* was possibly involved. The results showed that the number of microcolonies (MCs) was significantly reduced in the *LjPLP4* mutants, while the number of infection threads (ITs) that invaded to the epidermal cell increased compared to the wild type (Fig. S4a, b). The number of infection threads that branched from the tip and penetrated the cortical cells (Ramifying infection threads; Ramifying ITs) did not significantly increase overall (Fig. S4c). In addition, *LjPLP4* overexpression (*PLP4 OE*) also did not affect root growth (Fig. S5) but suppressed rhizobial infection without altering nodule number (Fig. 2e, f). These results strongly suggest that the function of *LjPLP4* is to suppress rhizobial infection in the epidermis. When *P. palmivora* was infected with the *L. japonicus* wild type and *LjPLP4* mutants, infection was suppressed in *Ljplp4* compared pathogen infection is opposite to that in rhizobial infection.

### LjPLP4 regulates rhizobial infection at the plasma membrane

LjPLP4 was shown to be localized to the PM in onion epidermal cells based on the observation using fluorescent protein (Kapranov *et al*., 2001). To investigate LjPLP4 subcellular localization in *L. japonicus*, Venus fluorescent protein fused to the N-terminal of LjPLP4 was expressed in roots. The Venus fluorescent pattern differed between the epidermal and cortical cells (Fig. 3a). In the cortical cells, dot-like structures were mainly observed at the periphery of the cells. In the epidermal cells, it appeared to be mainly localized at the PM, with a few dot-like structures similar to those in the cortical cells. The co-expression of the NF receptor NFR1 fused with mCherry, which was used as a PM marker, with Venus-LjPLP4 indicated that LjPLP4 was localized at the PM in root hairs (Fig. 3b, e) (Madsen *et al*., 2011). The dot-like structures were consistent with tdTomato-CTT, an endoplasmic reticulum (ER) marker (Fig. 3c, f) (Nagano et al., 2020). In addition, it was partly co-localized with a Golgi apparatus marker (Fig. 3d, g) (Ivanov & Harrison, 2014). These findings suggest that LjPLP4 functions differently between the epidermal and cortical cells based on its subcellular localization. To further specify the function of LjPLP4, the binding of LjPLP4 to phospholipids was examined using the PIP-strip assay. We used 6xHIS-LjPLP4 full-length (FL) or ΔSEC14 protein, in the latter the Sec14 domain was deleted, that was purified using the HIS tag (Fig. 3h). LjPLP4 FL bound to PI(3)P, PI(4)P, and PI(4,5)P_2_ but not to PI(3,5)P_2_ among phosphatidylinositol phosphates present in plants (Fig. 3i). To confirm the specificity and sensitivity of the LjPLP4 FL protein towards PI and phosphatidylinositol phosphates, array strips were used (Fig. 3j). These data indicated that LjPLP4 bound almost equally to PI(3)P, PI(4)P, and PI(4,5)P_2_, and these signals were lost when using ΔSEC14, indicating that binding was mediated by the SEC14 domain (Fig. 3k).

### LjPLP4 is not involved in Ca^2+^ spiking induced by NF recognition

To test whether LjPLP4, which has been suggested to function at the PM by subcellular localization analysis (Fig. 3a, b), affects the perception of NFs, we quantified Ca^2+^ spiking induced by NF treatment. We generated wild type and *Ljplp4* plants expressing Yellow Cameleon 2.60 (NLS-YC), a calcium indicator (Nagai *et al*., 2004; Soyano *et al*., 2024). NFs were added to the roots, and Ca^2+^ spiking was observed (Fig. S6). The results showed that the *LjPLP4* did not affect Ca^2+^ spiking because Ca^2+^ spiking occurred as frequently as in the wild type and no apparent differences in the spiking waveform observed (Fig. S6). This result suggests that LjPLP4 functions downstream of, or independent to, Ca^2+^ spiking induced by NFs.

### LjPIP5K4 and LjPIP5K6 are phosphatidylinositol 4-phosphate 5-kinase responsible for suppressing rhizobial infection

Among the three *LjPIP5Ks* whose expression levels were significantly upregulated during rhizobial infection (Fig. 1a), *LjPIP5K4* (LotjaGi1g1v0664200) and *LjPIP5K6* (LotjaGi3g1v0109700), encoding orthologous genes of *Arabidopsis PIP5K4* and *PIP5K6,* respectively, were selected for further investigation (Fig. S7). To confirm the tissue-specific expression of *LjPIP5K4* and *LjPIP5K6* during rhizobial infection, we performed GUS analysis with the promoter region of either *LjPIP5K4* or *LjPIP5K6*, and found that the GUS signal was induced by rhizobial infection in the epidermal cells where infection threads formed, and in the nodule primordia (Fig. 4a-d). We generated mutants of *LjPIP5K4* and *LjPIP5K6,* respectively, using CRISPR/Cas9, and confirmed that they did not affect root growth (Figs. S8 and S9). The loss of function in both of the genes resulted in an increase of infection threads compared with the wild type (Fig. 4e, g). The number of MCs was significantly reduced in the *LjPIP5K4* and *LjPIP5K6* mutants, while the number of Ramifying IT significantly increased compared to the wild type (Fig. S10). Subcellular localization analysis using the mCherry fluorescent protein suggested that LjPIP5K6 was co-localized with LjPLP4 at the PM in root hairs (Fig. S11). These results are consistent with the phenotype observed in the *Ljplp4* mutants, suggesting that LjPLP4 and LjPIP5K6 function concomitantly in the suppression of rhizobial infection. Furthermore, in both *Ljpip5k4* and *Ljpip5k6* mutants, the number of nodules was significantly higher than that in the wild type (Fig. 4f, h). The above results suggest that PI(4,5)P_2_, which is specifically produced by PIP5K, contributes to rhizobial infection in addition to nodule formation.

### *NIN* regulates the suppression of rhizobial infection partly through *LjPLP4*, *LjPIP5K4* and *LjPIP4K6*

Because overexpression of NIN in *L. japonicus* strongly suppresses the rhizobial infection (Soyano *et al*., 2014; Yoro et al., 2014), we investigated whether the suppression of rhizobial infection by *LjPLP4*, *LjPIP5K4* and *LjPIP4K6* is regulated by NIN. In the *nin-9* and *cyclops-6* mutant, the expression of *LjPLP4, LjPIP5K4* and *LjPIP5K*6 was not induced by inoculation of rhizobia (Fig. 5a-c), suggesting that the suppression of rhizobial infection by LjPLP4, LjPIP5K4 and LjPIP5K6 may be controlled by NIN and CYCLOPS. Therefore, to further investigate the relationship between *NIN* and *LjPLP4*, *LjPIP5K4* or *LjPIP5K6*, we generated *NIN*-overexpressing roots using the *L. japonicus* MG-20 wild type. In the NIN-overexpressing roots, the expression levels of *LjPLP4*, *LjPIP5K4*, and *LjPIP5K6* were found to be significantly higher compared to the roots expressing empty vector (EV) (Fig. 5d). Accordingly, the number of infection threads significantly increased in the *LjPLP4*, *LjPIP5K4*, and *LjPIP5K6* mutants compared to wild type plants overexpressing NIN, but not comparable to wild type plants expressing EV (Fig 5e). In *Ljpip5k4* and *Ljpip5k6* mutants, the suppression of nodule number by overexpression of *NIN* was weaker compared to the wild type, whereas in *Ljplp4*, significant difference was not observed compared to the wild type (Fig. 5f). The above results suggest that NIN regulates the suppression of rhizobial infection partly through *LjPLP4*, *LjPIP5K4* and *LjPIP4K6*. The suppression of rhizobial infection observed in *Ljplp4* led us to investigate whether *LjPLP4* is involved in the AON. It has been reported that the number of infection threads and nodules significantly increases in the AON mutants, *har1* and *tml* (Magori et al., 2009; Miri et al., 2019). A comparison of rhizobial infection phenotypes in *Ljplp4*, *har1-7*, and *tml* revealed that while there were significant differences in the number of nodules between *Ljplp4* and *har1-7* or *tml*, there were no significant differences in the number of infection threads (Fig. S12a, b). We quantified the *CLE-RS1* expression in *Ljplp4* roots 1 week after rhizobial inoculation and found that the expression level was induced similarly to the wild type but not in *nin-9*, indicating that the AON is triggered in *Ljplp4* roots (Fig. S12c). Furthermore, a grafting experiment between *Ljplp4* and the wild type was conducted to assess whether PLP4 was required in the shoot. It turned out that PLP4 functioned in the root to suppress infection thread formation (Fig. S12e,f). To know whether TML and/or HAR1 regulates *LjPLP4* expression, we examined *LjPLP4* expression in *tml* and *har1-7* roots 1 week after inoculation. The induction of *LjPLP4* expression by rhizobial infection that was observed in the wild type did not occur in *har1-7*, but it was significantly enhanced in *tml* (Fig. S12d). To further verify the relationship between the AON and *LjPLP4*, double mutants of *Ljplp4-4-10* and *har1-7* or *tml* were generated and were inoculated with rhizobia. The results showed that the number of infection threads significantly increased in *Ljplp4-4-10 har1-7* but not in *Ljplp4-4-10 tml*, when compared to *har1-7* or *tml* single mutants, respectively (Fig. 5g, i). There was no significant increase in the number of nodules when *Ljplp4* mutation was introduced in both *har1-7* and *tml*, consistent with the *Ljplp4* single mutant (Figs. 5h, j). These results suggest that the suppressive function of *LjPLP4* is partially HAR1-independent.

### Rhizobial infection-dependent PI(4,5)P_2_ accumulation is mediated by *LjPLP4* and *LjPIP5Ks*

The quantitative analysis using LC-MS showed that PI, PIP, and PIP_2_ accumulated in wild-type roots upon rhizobial infection (Fig. 1c-e). Therefore, we measured the total amounts of PI, PIP, and PIP_2_ in *Ljplp4* and *Ljpip5k6* mutants. The results showed that the amounts of PI, PIP, and PIP_2_ did not increase in *Ljplp4* and *Ljpip5k6* as observed in the wild type (Fig. S13). Analysis of *LjPIP5K4* and *LjPIP5K6* loss-of-function mutants suggested that PI(4,5)P_2_, among phosphatidylinositol phosphates, might have an important function in rhizobial infection (Fig. 4e, g). To examine specific changes in PI(4,5)P_2_ during rhizobial infection, we generated a transgenic *L. japonicus* MG-20 expressing 2xCHERRY-1xTUBBY-C (TUBBY), a fluorescent marker of PI(4,5)P_2_ (Simon *et al*., 2014). Strong signal was observed in the apex of root hairs (Fig. S14a). When roots were inoculated with *M. loti*, the signal was observed in the epidermis. In contrast, almost no signal was observed in the cortex (Fig. S14b). These findings indicate that accumulation of PI(4,5)P_2_ is induced in the root epidermis, where rhizobia enter host roots. Furthermore, when inoculated with *M. loti* carrying green fluorescent protein (GFP), PI(4,5)P_2_ accumulation was often found to be stronger in the epidermal cells where infection threads formed and in their adjacent cells, than in distal cells (Fig. 6a; Movie S1). Moreover, in the epidermal cells with stronger TUBBY signals, infection threads elongated horizontally often without defined direction (Fig. 6a), or became entangled and forming clumps (Fig. 6b). Infection threads in these cells seemed unable to penetrate the cortex from the epidermis. Interestingly, the TUBBY signal was shown to be much weaker where infection threads had penetrated cortex (Fig. 6c, d; Movie S2). The TUBBY signal in nodules was found to be absent where rhizobial infection had occurred and stronger in surrounding cells that were not infected with rhizobia (Fig. 6e). Because the number of infection threads increased in *Ljplp4*, *Ljpip5k4*, and *Ljpip5k6* mutants (Figs. 2b,c, 4e,g), we further verify the presence of PI(4,5)P_2_ in these mutants with TUBBY. The signal was almost absent in the cells where infection thread developed, indicating accumulation of PI(4,5)P_2_ was not induced by rhizobial infection (Fig. 6f-i).

**Figure 6.**
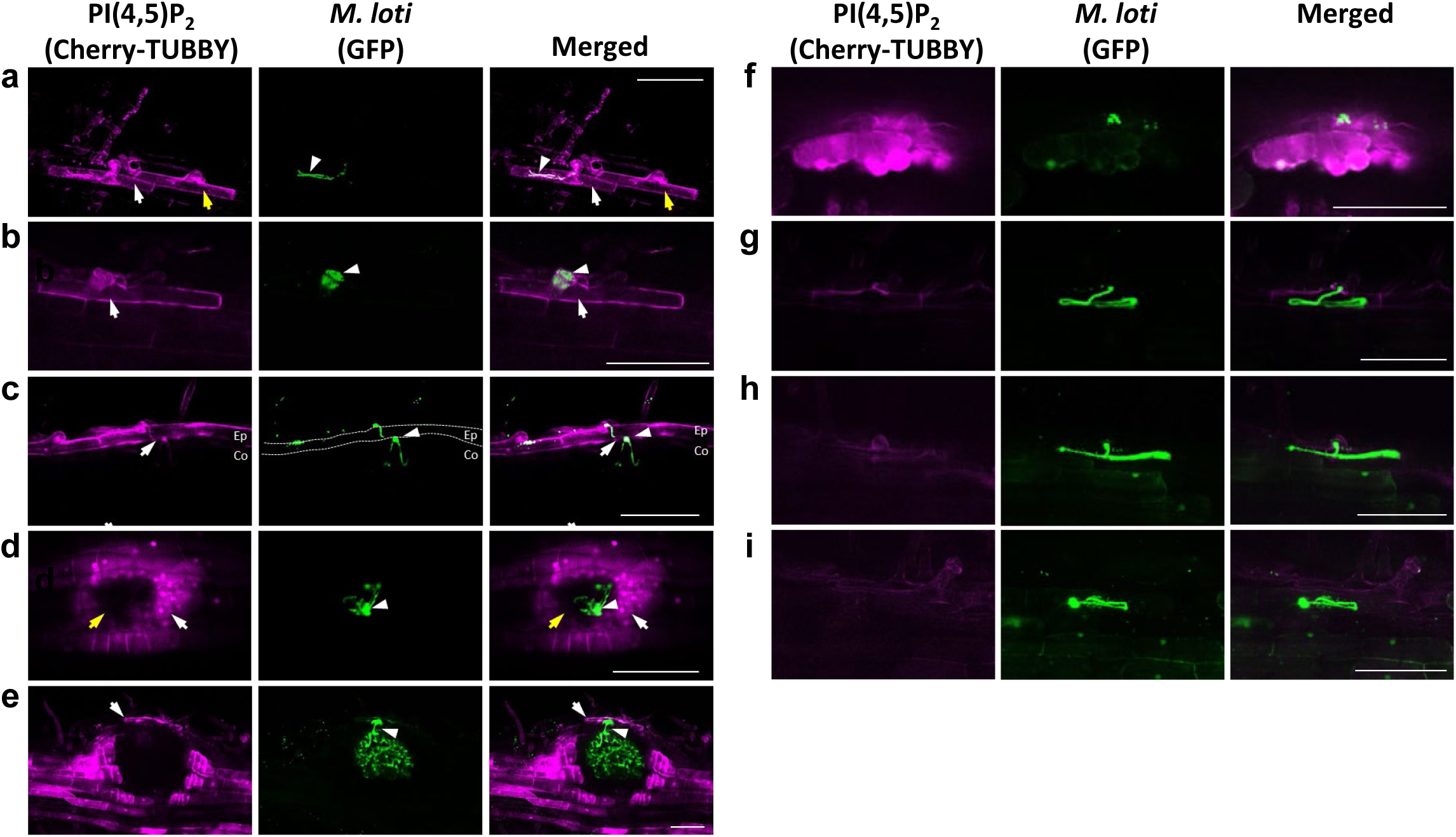
PI(4,5)P_2_ accumulation inhibits rhizobial invasion into root hair cells and elongation into the cortex. **a-e.** *L. japonicus* MG-20 wild-type expressing TUBBY, a PI(4,5)P_2_ marker, was infected with *M. loti* expressing GFP. The root epidermal cells (a,b), the epidermal and the cortex cells observed from horizontally (c), nodule primordium observed from above (d), immature nodule (e). **a.** White arrow indicates a TUBBY signal in an infected cell, yellow arrow indicates a TUBBY signal in the vicinity cells of infected cells, white arrowhead indicates infection thread that failed to penetrate to the cortex. **b.** White arrow indicates TUBBY signal in an infected cell; white arrowhead indicates infection thread that failed and forming clumps. **c.** White arrow indicates weakened accumulation of a TUBBY signal around infection threads penetrating to the cortex. Ep; Epidermis, Co; Cortex. **d.** White arrow indicates a TUBBY signal accumulated in the cells surrounding the infected region; Yellow arrow indicates a TUBBY signal is absent in the infected cell; The white arrowhead indicate infection threads that have branched off and are extending into cortex cell. **e.** White arrow indicates strong accumulation of a TUBBY signal on the surface of rhizobium-infected nodules. white arrowhead indicates infection threads invading cortical cells and rhizobia infecting cells inside the nodule. **f-i**. *L. japonicus* hairy roots expressing TUBBY with Venus as a transformation marker in wild type (f), *Ljplp4* (g), *Ljpip5k4* (h) or *Ljpip5k6* roots (i), were infected with *M. loti* expressing GFP. Scale bars is 100 µm (a-i).

## Discussion

In this study, RNA-seq analysis revealed rhizobial infection-dependent expression of PI signaling-related genes in *L. japonicus* roots (Fig. 1a). LC-MS analysis showed rhizobial infection-dependent increases in PI, PIP, and PIP_2_ (Fig. 1c–e). In the LC-MS analysis, quantifying individual phosphorylation variants in PIP and PIP_2_ was technically difficult. However, because there was no increase in total PIP_2_ in the *Ljpip5k6* mutant, PI(4,5)P_2_ likely accounts for much of the increase in PIP_2_ observed in these LC-MS data (Fig. S13) because PIP5K has been shown to specifically produce PI(4,5)P_2_ (Muftuoglu et al., 2016). In addition, observation of PI(4,5)P_2_ dynamics in rhizobia infected roots by TUBBY, a PI(4,5)P_2_ marker, revealed that ectopic PI(4,5)P_2_ accumulation was closely related to rhizobial infection (Fig. 6). Although we cannot rule out the possibility that PI and PIP function in rhizobial infection, the fact that increase of rhizobial infection not only in *LjPLP4* but also in *LjPIP5K4* and *LjPIP5K6* mutants (Figs. 2b, c and 4e, g) suggests that the shift from PI(4)P to PI(4,5)P_2_ has a suppressive effect on rhizobial infection.

Regarding the Sec14 domain present in LjPLP4, two functions have been postulated. First*, in vitro* lipid transport studies have shown that the Sec14 domain-containing protein transports PI and phosphatidylcholine (PC) across membranes (Kapranov *et al*., 2001; Schaaf *et al*., 2008). Second, in a model predicted based on the crystal structure, Sec14 binds to PI and PIPs on the membrane, thereby forming a structure that exposes PI and PIP to PI4K and PIP5K to be able to function efficiently (Kf de Campos & Schaaf, 2017). The real-time PCR and RNA-seq analyses performed in this study revealed that increased expression of *LjPLP4* 1 week after rhizobial inoculation (Fig. 1a, S1b). PIP strip assays revealed that LjPLP4 binds relatively strongly to PI(3)P, PI(4)P, and PI(4,5)P_2_ among PI and phosphatidylinositol phosphates present in plant cells (Fig. 3i, j). These findings also suggest that LjPLP4 facilitates PI(4)P and PI(4,5)P_2_ production by PI4K and PIP5K. This idea is consistent with the fact that the expression of one of *PI4K* genes (LotjaGi2g1v0442900) is significantly induced by rhizobial infection, along with three *LjPIP5Ks* (Fig. 1a). In addition, LjPLP4 is localized mainly at the PM and the endoplasmic reticulum (ER) and, to a lesser degree, to the Golgi bodies (Fig. 3a–g). The function of LjPLP4 in the ER and the Golgi bodies remains unclear, but as LjPLP4 has been shown to be localized at PM in the root epidermal cells, which are the main contact tissue with soil microorganisms, it probably functions to synthesize PI(4)P and PI(4,5)P_2_. Observations of PI (4,5)P_2_ by TUBBY showed a hardly detectable signal in the cortex (Fig. S13b). This may be related to the fact that *LjPLP4* is not PM-localized in the cortex (Fig. 3a, b). The expression pattern analysis using GUS staining revealed that *LjPLP4* was expressed in the epidermis and the cortex in inoculated roots, while *LjPIP5K4* and *LjPIP5K6* were expressed specifically in rhizoial infected cells. The differential expression pattern suggests that LjPLP4 is involved not only with LjPIP5K4 and LjPIP5K6 but also with other PIP5Ks. *LjPIP5K4* and *LjPIP5K6* were expressed in the nodule primordium. The results suggest that *LjPIP5K4* and *LjPIP5K6* may be involved in nodule development. However, the increased number of nodules observed in the *Ljpip5k4* and *Ljpip5k6* mutants appears to result from an increased number of ramifying infection threads in these mutants. These results seem to conflict with the finding that PI(4,5)P_2_ was absent in nodule primordium (Fig. 6e). It has been reported that diacylglycerol (DAG) accumulates in the nodules in *M. truncatula* (Zhang et al., 2020). Because PI(4,5)P_2_ is hydrolyzed by phospholipase C (PLC) to produce DAG and IP_3_, the PI(4,5)P_2_ generated by LjPIP5K and LjPIP5K6 may contribute to nodule formation by being converted into DAG and/or IP_3_.

In animal cells, it is known that PI(4, 5)P_2_ cleavage by phospholipase C (PLC) releases inositol 1,4,5-trisphosphate (IP_3_), which functions as a second messenger to trigger Ca^2+^ channel opening (Kania *et al*., 2017). Furthermore, in *M. truncatula*, the IP_3_ inducer Mastoparan analog (Mas7) treatment induces calcium fluctuations similar to NF-induced Ca^2+^ spiking (Sun *et al*., 2007). In this regard, we investigated whether the absence of *LjPLP4* affected Ca^2+^ spiking induced by NFs using a YC 2.60 sensor but found no significant variation in the spiking (Fig. S6). Thus, it is unlikely that Ca^2+^ spiking is affected by *LjPLP4*-induced changes in PI dynamics.

The *NIN*-dependent suppression mechanism of rhizobial infection has been demonstrated previously (Soyano *et al*., 2014; Yoro *et al*., 2020). The real-time PCR analysis with the *nin-9* mutant found that the expression of *LjPLP4*, *LjPIP5K4*, and *LjPIP5K6* required *NIN* (Fig. 5a-c). In addition, in the *LjPLP4*, *LjPIP5K4*, and *LjPIP5K6* mutants overexpressing *NIN*, the number of infection threads was significantly higher than in wild-type plants overexpressing *NIN* (Fig. 5e). The above results strongly suggest that the suppression of rhizobial infection by *LjPLP4*, *LjPIP5K4*, and *LjPIP5K6* is partly regulated by *NIN*.

Expression analysis in this study showed that the *Ljplp4* mutation did not affect the expression of *CLE-RS1* (Fig. S12c). NIN regulates expression levels of downstream genes, including *CLE-RS1*, in a rhizobial symbiosis-specific manner (Soyano *et al*., 2014). The *Ljplp4 har1* double mutant showed an increase in the number of infection threads compared with the *Ljplp4* single mutant (Fig. 5i). In the *har1-7* mutant, *NIN* is expressed even in the absence of rhizobial infection (Akamatsu et al., 2021). Therefore, the gene expression analysis in *har1-7* may not have detected an increase in *PLP4* expression (Fig. S12d). The results suggest that the suppression of rhizobial infection mediated by *LjPLP4* is likely independent of *HAR1*. In the *Ljplp4 tml* double mutant, no increase in the number of infection threads was observed compared to the *tml* single mutant (Fig. 5e). Gene expression analysis found that *LjPLP4* was induced in a *TML*-independent manner (Fig. S12d). The above results suggest that while *LjPLP4* is *TML*-independent in terms of gene expression, LjPLP4 has a functional connection with TML in the suppression of rhizobial infection.

PI(4,5)P_2_ has been shown to accumulate at the tip of infection threads in the epidermis in *M. truncatula* (Lace et al., 2023). Previous studies have suggested that PI(4,5)P_2_ plays a role in regulating the cytoskeleton and pectin by accumulating at sites of tip growth (Ischebeck *et al*., 2008; Hirano *et al*., 2018). Thus, PI(4,5)P_2_ may promote elongation of infection threads in root hairs. On the other hand, in this study, we showed that in *L. japonicus*, the epidermal cells in the wild type in which infection threads are aborted show the clear accumulation of PI(4,5)P_2_ to the PM (Fig. 6a, b). In addition, in *LjPLP4*, *LjPIP5K4* and *LjPIP5K6* mutants, the accumulation of PI(4,5)P_2_ in the epidermis were weaker than in the wild type (Fig. 6f-i). Moreover, analysis of the rhizobial infection phenotype in *LjPLP4*, *LjPIP5K4* and *LjPIP5K6* mutants suggests that PI(4,5)P_2_ negatively regulates rhizobial infection (Figs. 2b, c, 4e, g). The above results suggest that accumulation of PI(4,5)P_2_ to the PM likely facilitates the suppression of rhizobial infection. In *M. truncatula*, the infection chamber formed at the tip of root hairs during rhizobial infection becomes enlarged as the first step of infection thread formation (Furnier *et al*., 2015). The enlarged membrane is then polarized by a protein complex containing RPG, VPY and LIN and proceeds to invade in the direction of the root center. PI(4,5)P_2_ accumulates at the tip of the polarized infection thread, which is crucial for the elongation of infection thread with correct direction (Liu et al., 2019; Lace et al., 2023). Based on this, it is hypothesized that the strong accumulation of PI(4,5)P_2_ to the PM causes the loss of polarity, preventing the penetration of infection threads into the cortex. Furthermore, quantitative analysis of infection events revealed a decrease in the number of MCs in the *LjPLP4*, *LjPIP5k4*, and *LjPIP5K6* mutants, suggesting that the PI(4,5)P_2_ accumulation to the PM inhibited the entry of rhizobia into the root hair cells. How PI(4,5)P_2_ accumulates to the PM and suppresses excessive rhizobial infection is currently unknown. One possibility is that excess PI, PIP, and PIP_2_ are negatively charged and thereby may impair the recruitment of factors necessary for rhizobial infection, inhibiting rhizobial infection. These questions must be investigated in future research studies.

In host plant immune responses to pathogenic fungi and oomycetes, the importance of PI(4,5)P_2_ or PI(4)P at the infection site is becoming clear (Shimada *et al*., 2019; Qin *et al*., 2020; Zarreen *et al*., 2023). As the local accumulation likely promotes pathogen infection, it has been thought that pathogens can regulate PI(4,5)P_2_ production and/or localization in host cells by contacting their invasive hyphae (Shimada *et al*., 2019). Although no upregulation of *LjPLP4* was observed during *P. palmivora* infection, the lesion length of *P. palmivora* infection, and *P. palmivora* RNA content were markedly suppressed in the *LjPLP4* mutant, suggesting that *LjPLP4* plays a positive role in *P. palmivora*infection (Fig. 1b, 2g–i). The gene expression of *PI4Ks* (LotjaGi3g1g1v0712000_LC, LotjaGi3g0396100) was induced in *P. palmivora* infection, but not in rhizobial infection (Fig. 1b), suggesting that these PI4K might function together with LjPLP4 in defense responses. These results indicate that a similar function to ectopic PI(4,5)P_2_ accumulation at the EHM observed during powdery mildew infection reported in *Arabidopsis* is also present in *L. japonicus* and that it requires *LjPLP4*.

In summary (Fig. 7), PI signaling-related genes are upregulated in rhizobial-infected *L. japonicus* roots. Among them, *LjPLP4* causes PI(4,5)P_2_ accumulation in the root epidermis. The accumulation of PI(4,5)P_2_ in the epidermis by inoculation of rhizobia inhibits the invasion of rhizobia into root hairs. The expression of *LjPLP4*, *LjPIP5k4* and *LjPIP5k6* regulated by *NIN* suggests that these responses are likely to be rhizobial infection-specific phenomena. Although the direct interaction of LjPLP4 and LjPIP5Ks has not been demonstrated in this study, co-localization and requirement for accumulation of PI(4,5)P_2_ suggest that they may function together in the epidermal cells where infection threads formed. In the cells where PI(4,5)P_2_ does not accumulate or is degraded, rhizobia invade into root hairs. Furthermore, inhibition due to increased PI(4,5)P_2_ also affects elongation of infection threads. Only a few infection threads that avoid the accumulation of PI(4,5)P_2_ are able to proceed with infection into the cortex.

**Figure 7.**
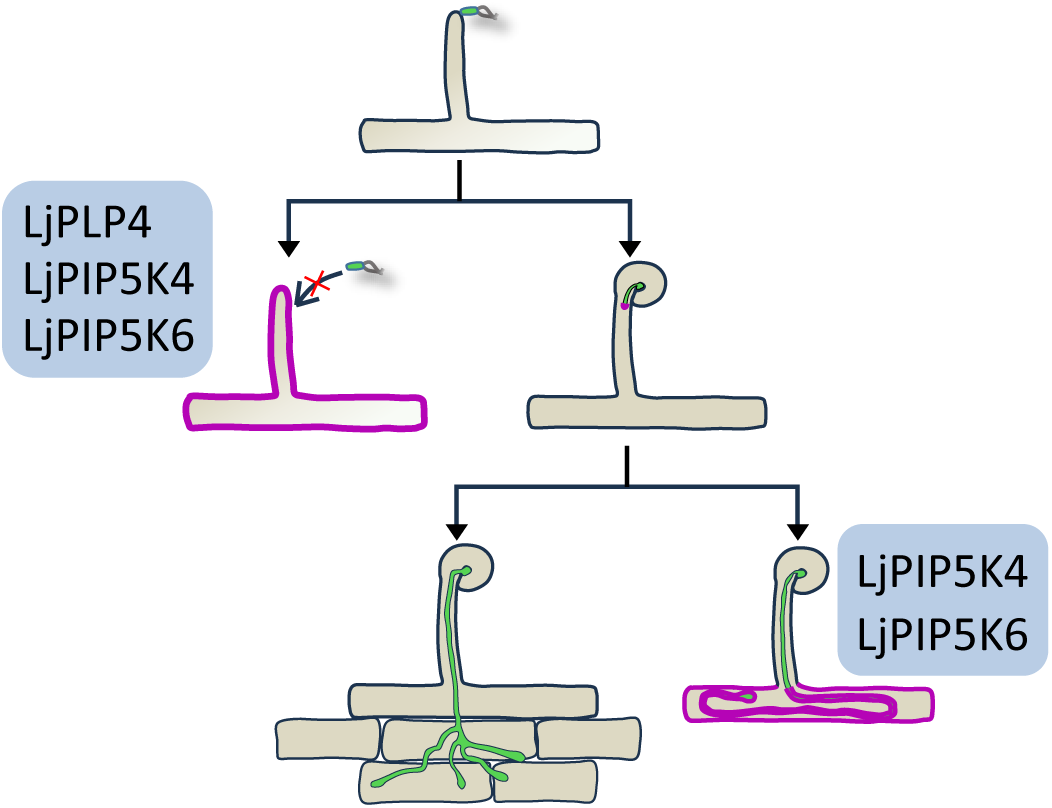
Schematic model of the pathway of the suppression system by PI(4,5)P_2_. The accumulation of PI(4,5)P_2_ in the epidermis by inoculation of rhizobia inhibits the invasion of rhizobia into the root hairs. In the cells where PI(4,5)P_2_ does not accumulate or is degraded, rhizobia invade into the root hairs. Furthermore, the accumulation of PI(4,5)P_2_ also affects elongation of infection threads with correct direction. Only a few infection threads that avoid accumulation of PI(4,5)P_2_ are able to penetrate the cortex. The green color indicates infection thread, and the magenta color indicates PI(4,5)P_2_.

## ACKNOWLEDGEMENTS

This work was supported by JSPS KAKENHI Grant Numbers 20K15426 22K06288 (to A.A) 23K17998 (to N. T), an Individual Special Research Fund (Kwansei Gakuin University, Japan, 2023), Hyogo Science and Technology, the Sumitomo Foundation and the Joint Usage/Research Center, the Institute of Plant Science and Resources (Okayama University, Japan). The CRISPR/Cas9 vectors were kindly provided by Dr. Masaki Endo (NARO, Japan). We thank Dr. Masayoshi Kawaguchi (National Institute for Basic Biology, Japan) and Dr. Takuya Suzaki (University of Tsukuba) for kindly providing *nin-9*, *tml*, *har1-7* and *cyclops-6* mutants. We thank Dr. Ayaka Hieno for the lecture on *P. palmivora* culture methods. We thank Mr. Hideki Nishimura for his technical support for *P. palmivora* culture. We acknowledge the NGS core facility of the Genome Information Research Center at the Research Institute for Microbial Diseases of Osaka University for the support in RNA sequencing and data analysis. pCMB-GAr was a gift from Maria Harrison (Addgene plasmid # 61173 ; http://n2t.net/addgene:61173 ; RRID:Addgene_61173) and tdTomato-CTT was a gift from Dr. Minoru Nagano (Ritsumeikan University).

## Author contributions

A.A designed the study. A.A, M.H and N.T wrote the manuscript. T.I contributed to the PIs measurements. H.T and Y.K contributed to data collection. All authors contributed to the interpretation of data and reviewed the manuscript.

## DATA AVAILABILITY STATEMENT

RNA-Seq data used in this publication have been deposited in DDBJ Sequence Read Archive (DRA) at the DNA Data Bank of Japan (DDBJ; http://www.ddbj.nig.ac.jp/) under the accession number DRR585568-DRR585579.

