## Supplementary Materials for "Rhizobial infection-specific accumulation of phosphatidylinositol 4,5-bisphosphate inhibits the excessive infection of rhizobia in *Lotus japonicus*"

### Supplementary Figure 1

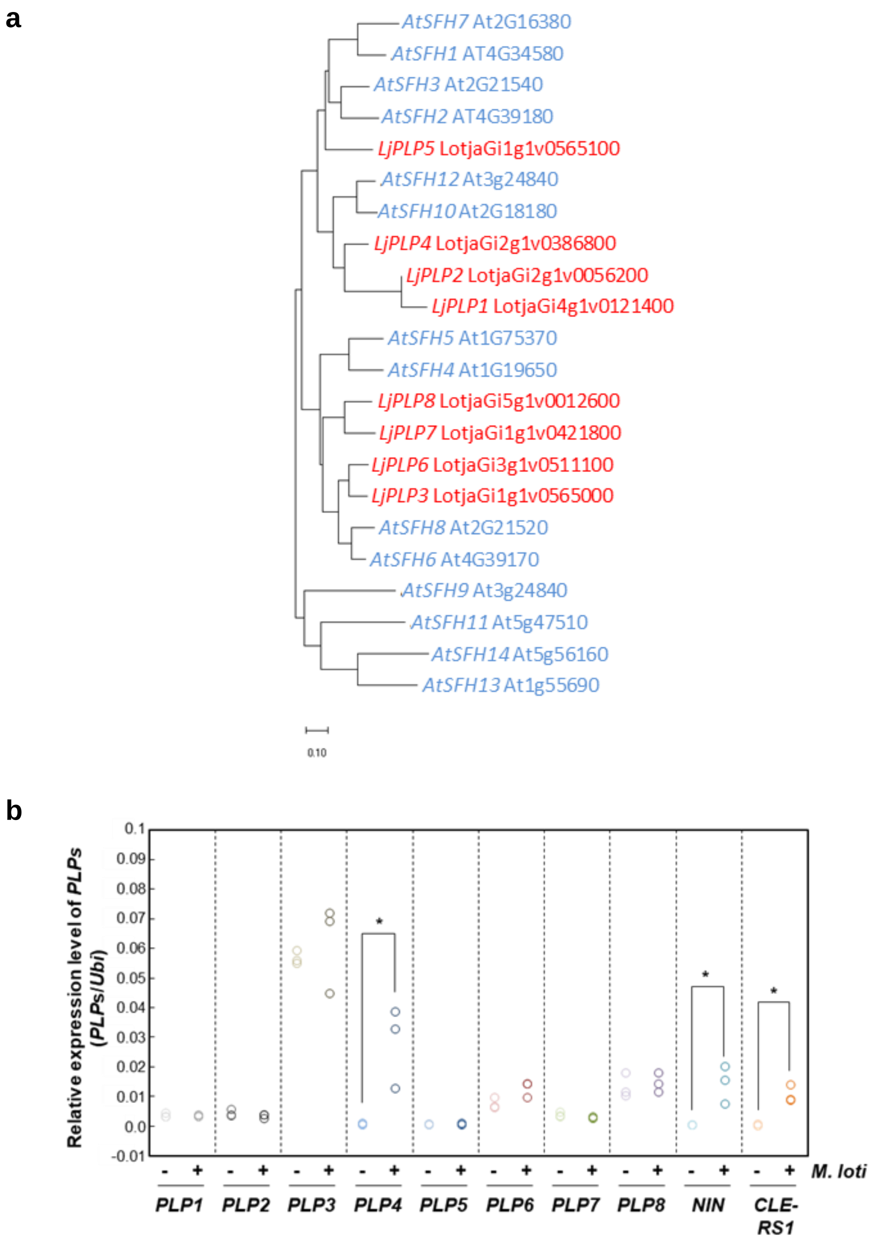


**Supplementary Figure 1 a.** Phylogenetic tree was constructed with MEGA X software. There are 8 Sec14-Nodulin genes in *Lotus japonicus* MG-20 and 14 SEC14-nodulin genes in *Arabidopsis thaliana*. **b**. Real-time PCR analysis of mRNA expression levels of SEC14-Nodulin genes in *L. japonicus* roots inoculated with *Mesorhizobium loti* for 1 week. (n = 3, *t*-test, *P< 0.05)

### Supplementary Figure 2

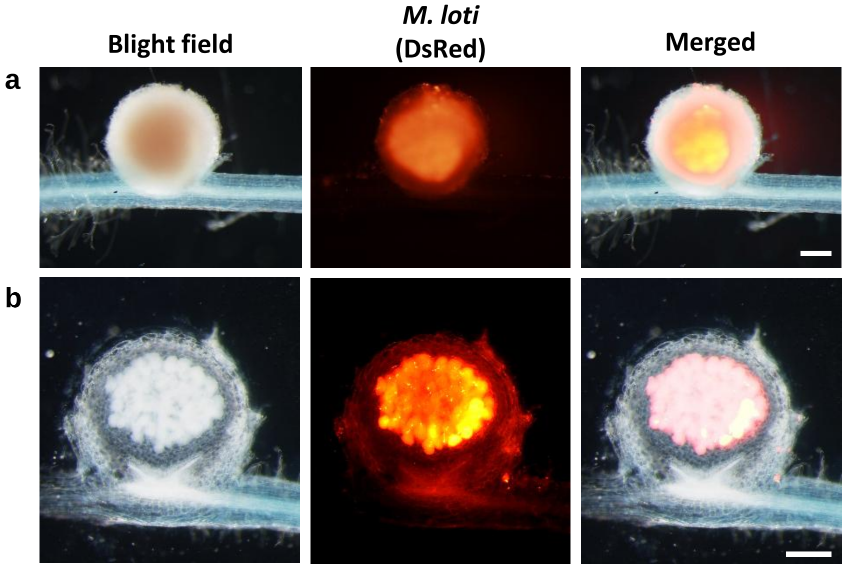


**Supplementary Figure 2** Rhizobia-infected *L. japonicus* MG-20 wild-type (WT) roots transformed with *pLjPLP4::GUS*. An infected nodule (a), 100 µm section of an infected nodule (b). The roots were stained with GUS staining buffer at 2 wai of *M. loti* carrying DsRed. Scale bars, 200 µm.

### Supplementary Figure 3

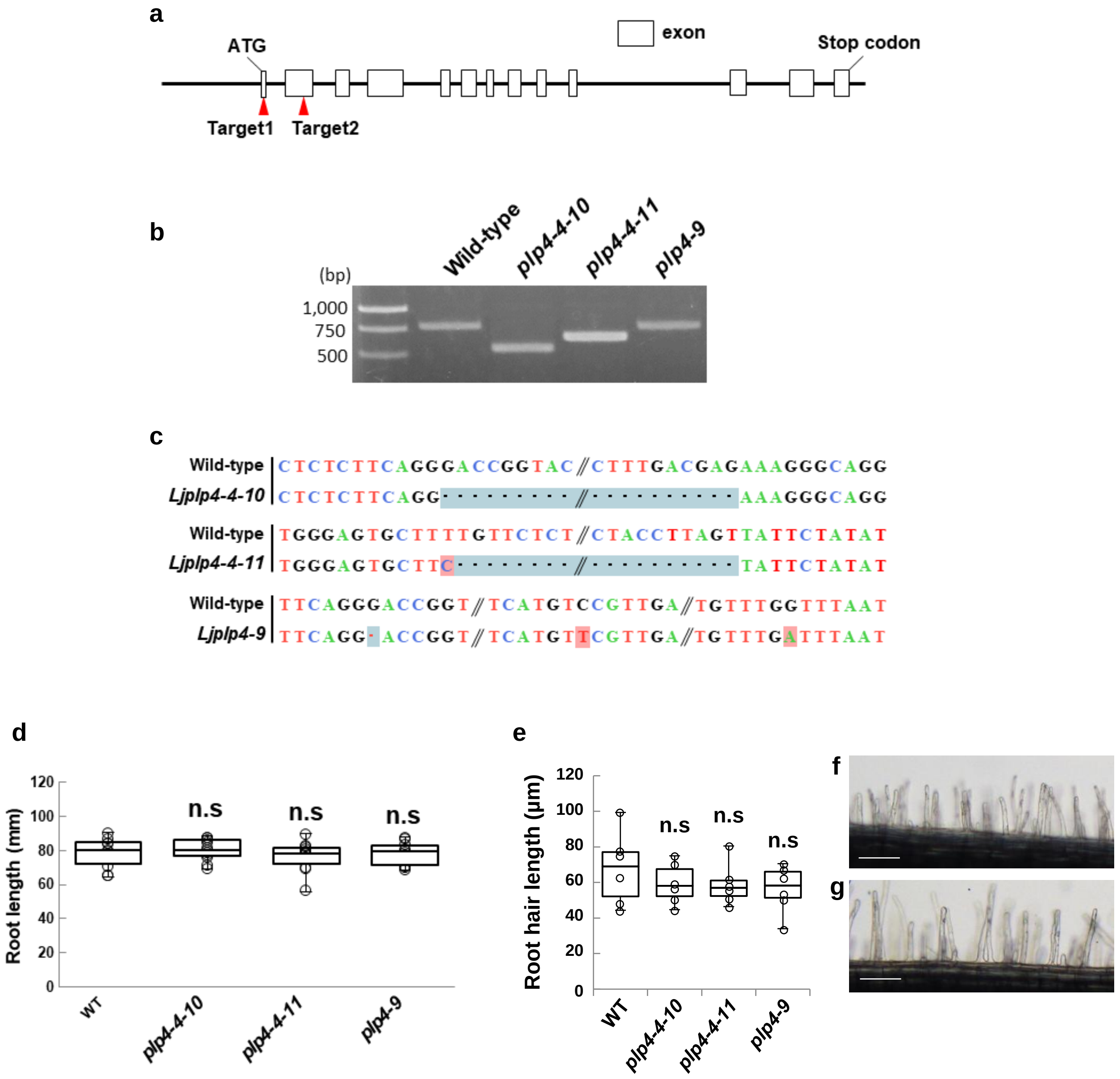


**Supplementary Figure 3**. Characterization of deletion mutants of *LjPIP4* using the CRISPR/Cas9 system. **a**. The genomic sequence of *LjPLP4* with CRISPR/Cas9 target sites. **b**. The *LjPLP4* fragment, including two CRISPR/Cas9 targets, was amplified from wild type plants (774 bp) and three mutants by PCR. **c**. Sequences surrounding the deletion or mutation site compared with the wild type sequence. *plp4-4-10*: 63 bp deletion including 1st exon and 1st intron, *plp4-4-11*: deletion from 5'UTR to 1st exon including start codon, *plp4-9*: 1 bp deletion at Target1 site and two point mutations. **d.** Root length at 1 wai with *M. loti*. Dunnett's test, $P<0.05$, n.s = not significant, $n\geq7$. **e-g**. Analysis of root hair growth. **e.** Root hair length at 2 days after germination. Dunnett's test, $P<0.05$, $n=6$. **f, g.** Root hairs observed in wild type (f) and *Ljplp4-4-10* (g). Scale bars, 100 μm.

### Supplementary Figure 4

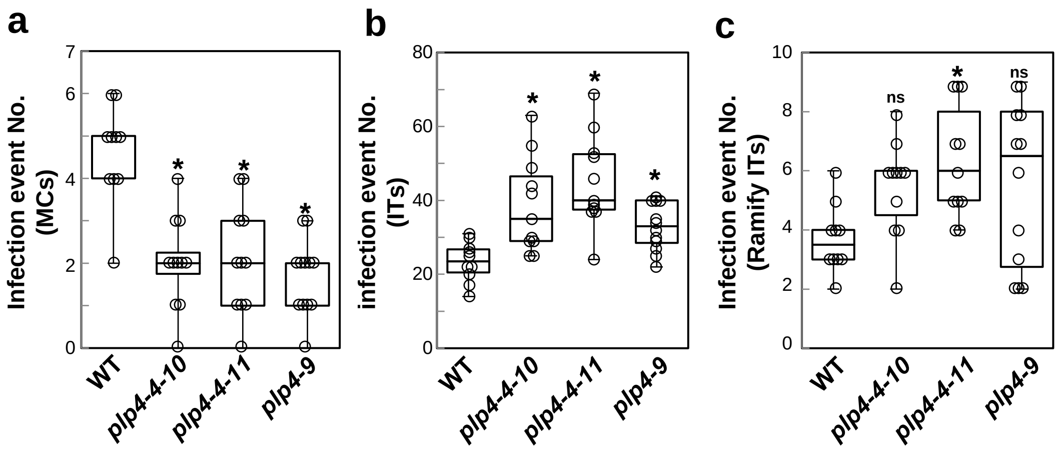


**Supplementary Figure 4** Quantitative analysis infection events. *L. japonicus* MG-20 wild-type (WT), *Ljplp4-4-10*, *Ljplp4-4-11* and *Ljplp4-9* plants were inoculated with *M. loti* carrying DsRed for 7 days. MCs; Microcolonies, ITs; infection threads that penetrated to the epidermal cells, Ramify ITs; infection threads that branched from the tip and penetrated the cortical cells. MCs (a), ITs (b), and Ramifying ITs (c) were counted. n ≥ 10, Dunnett's test, one asterisk (*) indicates $P < 0.05$, two (**) $P < 0.01$.

### Supplementary Figure 5

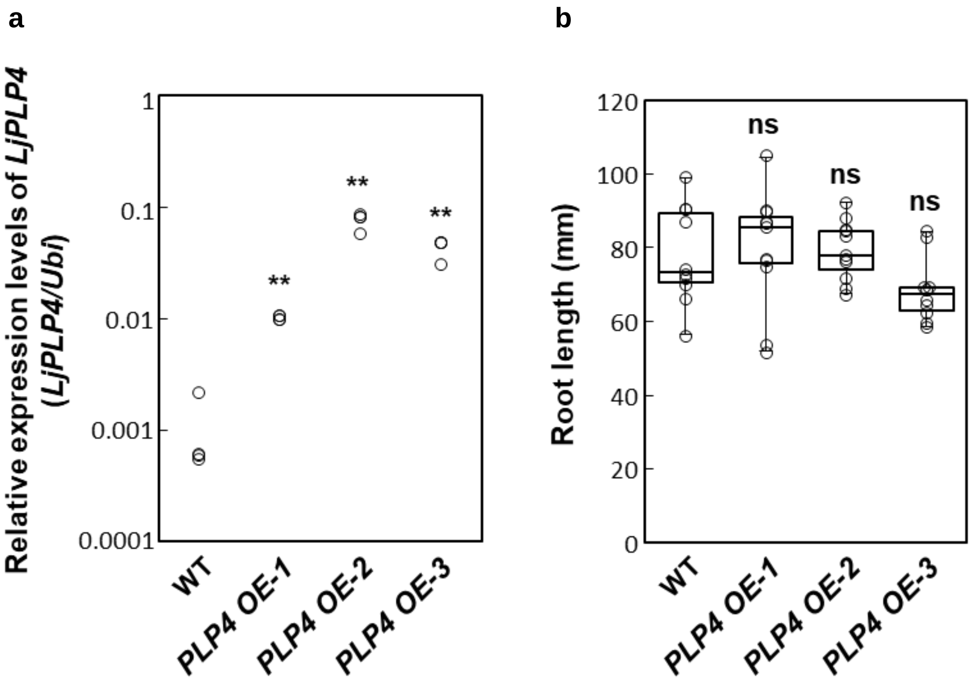


**Supplementary Figure 5**. Characterization of overexpression plants of *LjPLP4*. **a**. real-time PCR of *LjPLP4* in overexpression (OE) plants driven by *L. japonicus* Ubiquitin promoter. *PLP4 OE-1*, *PLP4 OE-2* and *PLP4 OE-3* are independent LjPLP4 OE lines. Relative expression compared with Ubiquitin expression. **b.** Boxplots of root length at 1 wai with *M. loti* carrying DsRed. Dunnett's test, **P<0.01, ns = not significant, n = 4 (a), n ≥ 7(b).

### Supplementary Figure 6

**a**

| | Spiking (+) | Total cells | % [(+)/Total cells] | No. of plants |
|---|---|---|---|---|
| MG20 wild type | 64 | 90 | 71.1 | 9 |
| *plp4-4-10* | 93 | 120 | 77.5 ns | 12 |

**b**

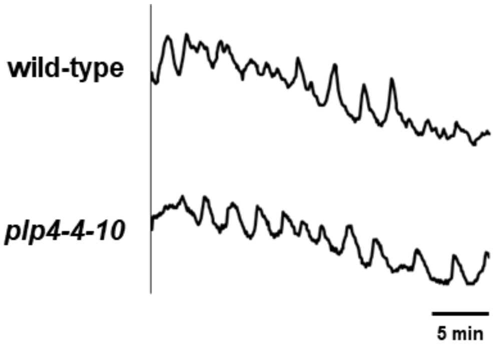


**Supplementary Figure 6**. Observation of the $Ca^{2+}$ spiking in *L. japonicus* MG-20 wild type and *Ljplp4-4-10* in the presence of $10^{-8}$ M of Nod factors. **a.** Percentage of the number of cells with $Ca^{2+}$ spiking relative to the number of cells observed. (chi-square test, ns indicate no significant differences). **b.** Typical $Ca^{2+}$ spiking patterns induced Nod factor treatment in wild-type and *Ljplp4-4-10* mutant.

### Supplementary Figure 7

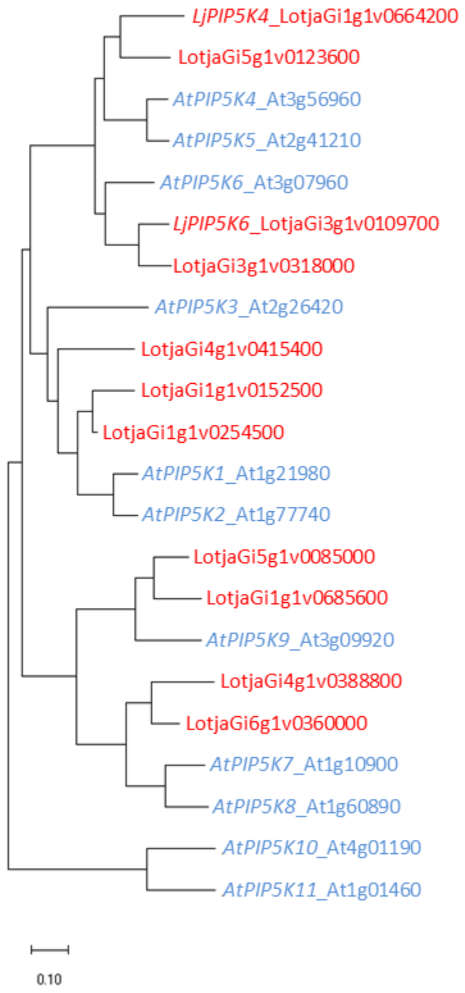


**Supplementary Figure 7** Phylogenetic tree was constructed with MEGA X software. There are 11 PIP5K genes in *Lotus japonicus* MG-20 and 11 genes in *Arabidopsis thaliana*.

### Supplementary Figure 8

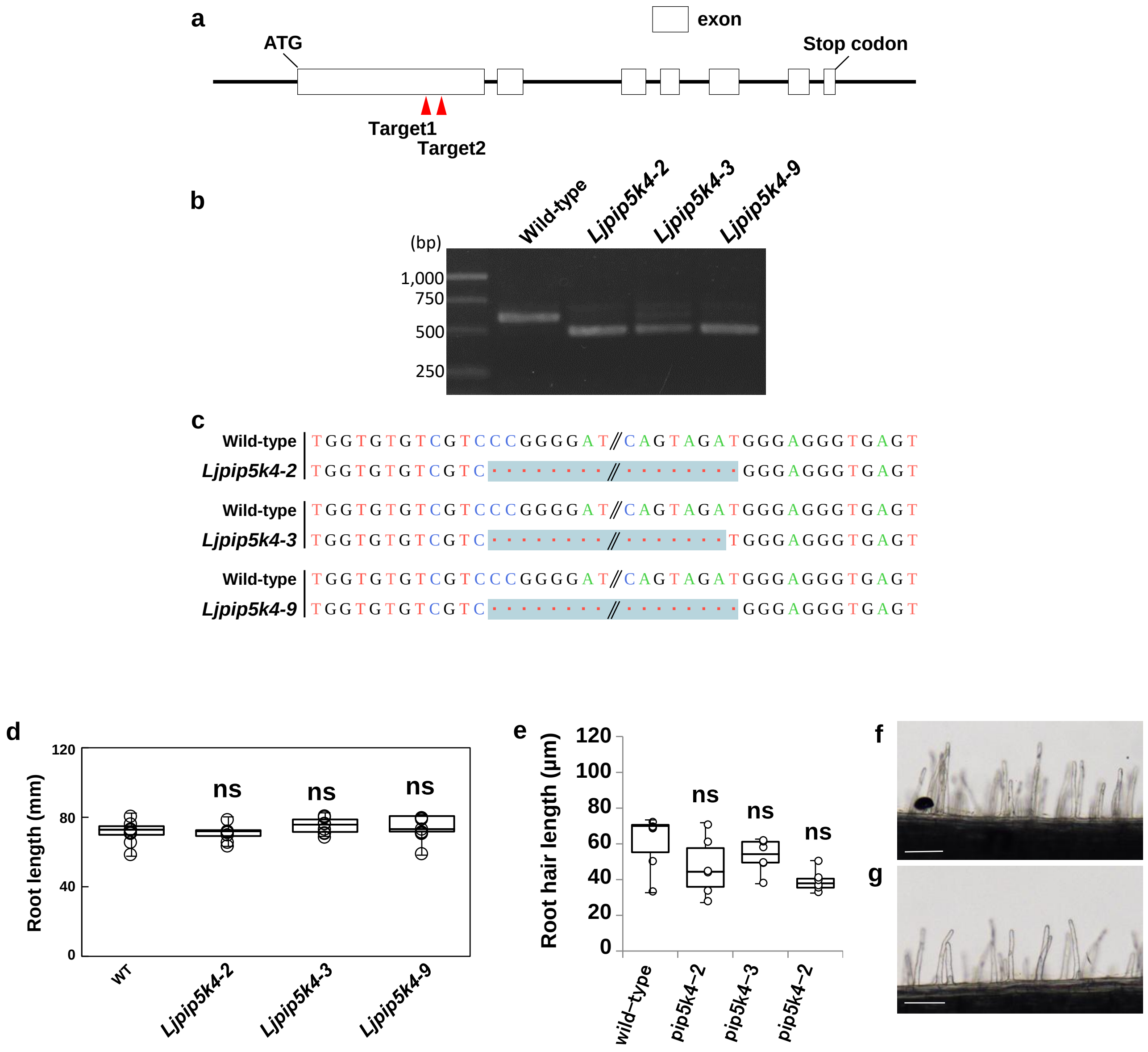


**Supplementary Figure 8**. Characterization of deletion mutants of *LjPIP5K4* using CRISPR/Cas9 system. **a**. The genomic sequence of *LjPIP5K4* with CRISPR/Cas9 target sites. **b**. The *LjPIP5K4* fragment, including two CRISPR/Cas9 targets (600 bp), was amplified from wild-type plants and three mutants by PCR. **c**. Sequences surrounding the deletion or mutation site compared with the wild-type sequence. *Ljpip5k4-2*: 101 bp deletion from target site1 to target site2, p*ip5k4-3*: 100 bp deletion from target site1 to target site2, p*ip5k4-9*: 101 bp deletion from target site1 to target site2. **d.** Boxplots of root length at 1 wai with *M. loti* carrying *DsRed*. (Dunnett's test, P<0.05, ns = not significant, n ≥ 7). Similar results were obtained in three independent experiments. **e-g**. Analysis of root hair growth. **e.** Boxplots of root hair length at 2 days after germination (Dunnett's test, P<0.05, n = 6). **f, g.** Root hairs observed in wild type (f) and *Ljpip5k4-2* (g). Scale bars, 100 µm.

### Supplementary Figure 9

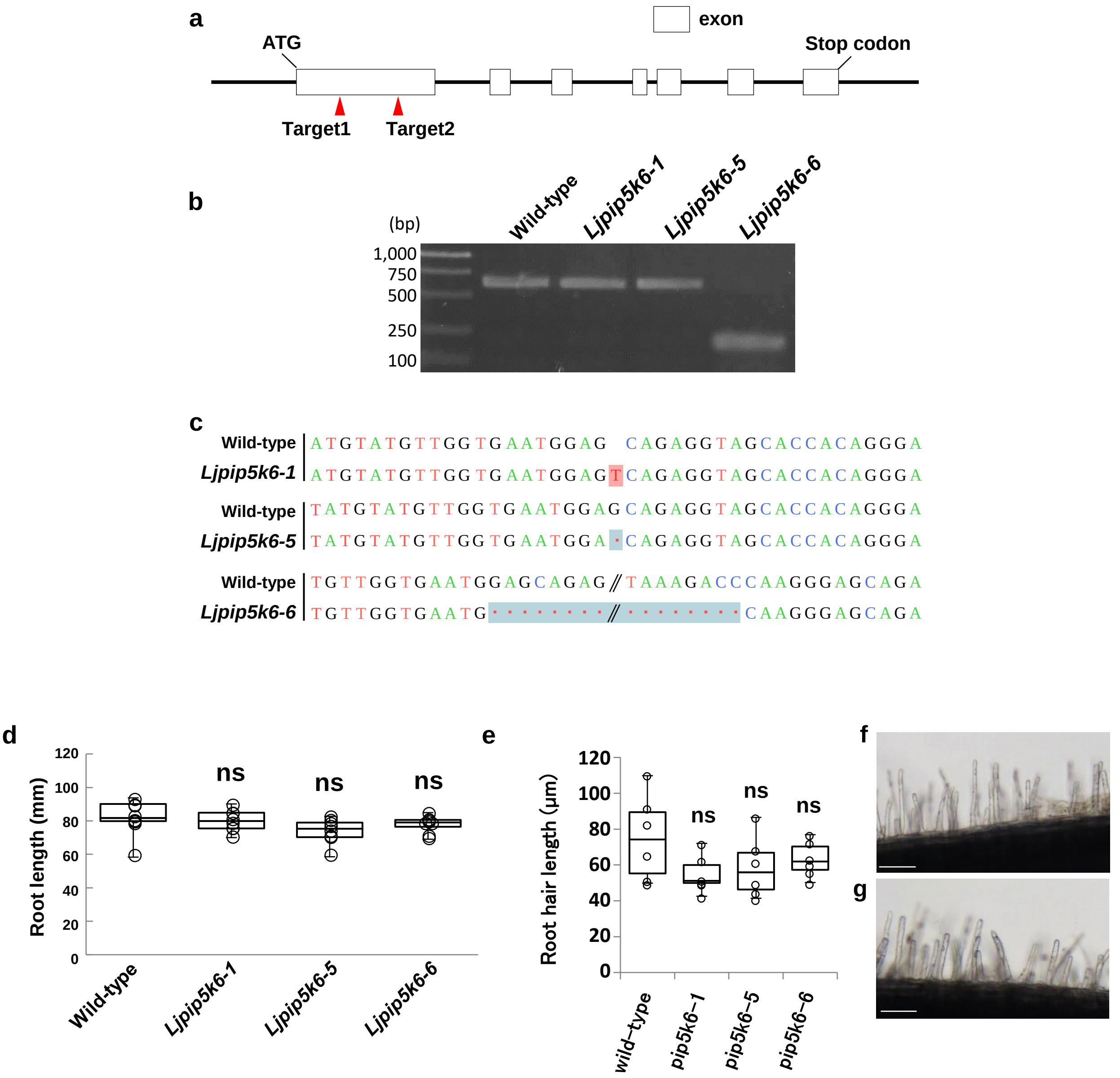


**Supplementary Figure 9**. Characterization of deletion mutants of *LjPIP5K6* using CRISPR/Cas9 system. **a**. The genomic sequence of *LjPIP5K6* with CRISPR/Cas9 target sites. **b**. The *LjPIP5K6* fragment, including two CRISPR/Cas9 targets (614 bp), was amplified from wild-type plants and three mutants by PCR. **c**. Sequences surrounding the deletion or mutation site compared with the wild-type sequence. *pip5k6-1*: 1 bp insertion at target1 site, *pip5k6-5*: 1 bp deletion at target1 site, *pip5k-6*: a deletion in first exon from target site1 to target site2. **d.** Boxplots of root length at 1 wai with *M. loti* carrying *DsRed*. Dunnett's test, $P<0.05$, ns = not significant, $n \geq 7$. Similar results were obtained in three independent experiments. **e-g**. Analysis of root hair growth. **e.** Boxplots of root hair length at 2 days after germination. Dunnett's test, $P<0.05$, $n = 6$. **f, g.** Root hairs observed in wild type (f) and *Ljpip5k6-1* (g). Scale bars, 100 μm.

### Supplementary Figure 10

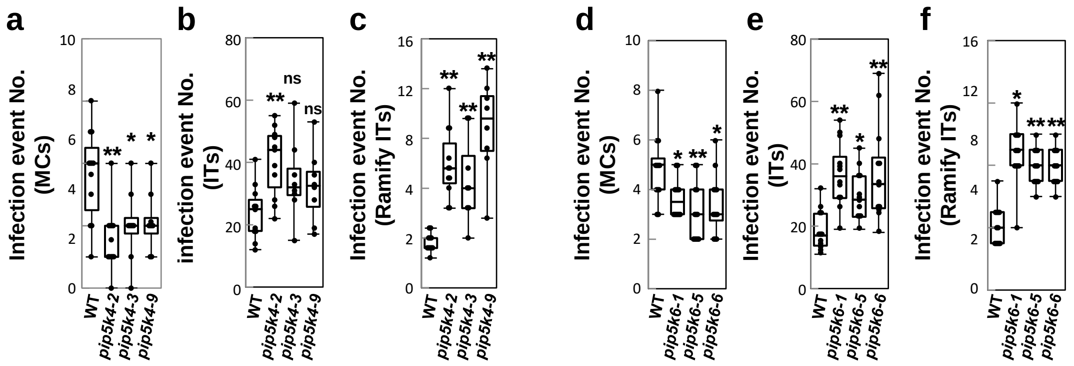


**Supplementary Figure 10** Quantitative analysis of infection events. *L. japonicus* MG-20 wild-type (WT), *Ljplp4-4-10*, *Ljplp4-4-11* and *Ljplp4-9* plants were inoculated with *M. loti* carrying DsRed for 7 days. Microcolonies [(MCs; (a, d)], infection threads (ITs) that invaded to the epidermal cells (b, e), and infection threads that branched from the tip and penetrated the cortex [Ramifying infection threads; Ramifying ITs; (c, f)] were counted. n ≥ 10, Dunnett's test, one asterisk (*) indicates $P < 0.05$, two (**) $P < 0.01$.

### Supplementary Figure 11

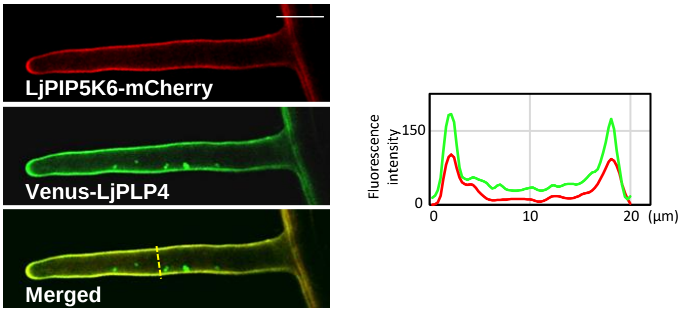


**Supplementary Figure 11**. The plasma membrane localization of LjPIP5K6 in *L. japonicus* roots**.** *L. japonicus* wild-type hairy roots transformed with fluorescent constructs *LjPIP5K6-mCherry* (Red) and *Venus-LjPLP4* (green). Scale bars is 20 µm.

### Supplementary Figure 12

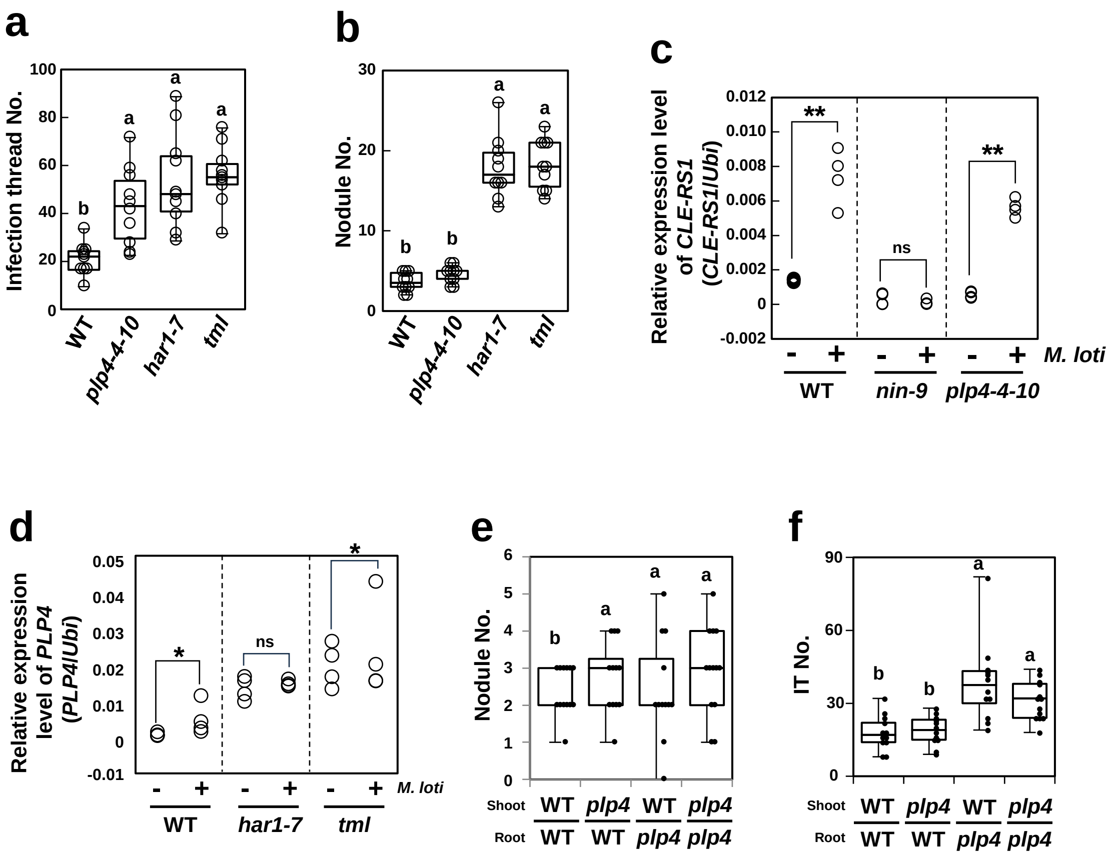


**Supplementary Figure 12** The plants were inoculated with *M. loti* carrying *DsRed* (a, b) or wild-type (c, d) for 1 week. **a, b.** Rhizobial infection phenotype in *L. japonicus* wild-type, *Ljplp4-4-10*, *har1-7*, and *tml* mutants. **c, d.** Quantitative analysis of *CLE-RS1* expression in *nin-9* and *Ljplp4-4-10* mutants (c), *LjPLP4* expression in wild-type, *har1-7,* and *tml* (d) by real-time PCR. [n ≥ 8, Tukey test, *P< 0.05, (a, b), n = 4, *t*-test, **P < 0.01. *P < 0.05 (c, d)], **e, f.** Shoot-root grafting with wild type and *Ljplp4-4-10*. Grafted plants were inoculated with *M. loti* carrying *DsRed*. Nodules and infection threads per plant were counted at 7 dpi. The shoot and the root genotype used for each grafting combination are shown. n ≥ 12, Tukey test, *P< 0.05.

### Supplementary Figure 13

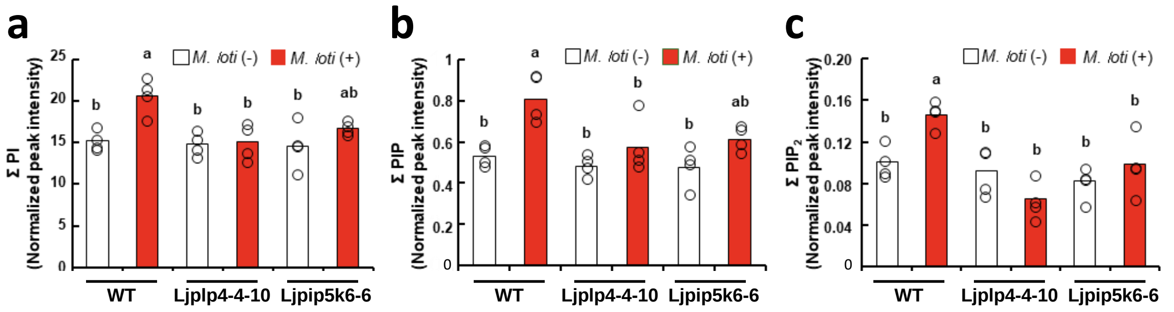


**Supplementary Figure 13**. LC-MS analysis of PI (a), PIP (b), and $PIP_2$ (c) in *L. japonicus* MG-20 roots of wild-type, *Ljplp4-4-10,* and *Ljpip5k6-6*, in control conditions (white) or inoculated with *M. loti* for 1 weed (Red) (n = 4 biological replicates, the bars show average, and circles show each data point, Tukey test; $*P < 0.05$).

### Supplementary Figure 14

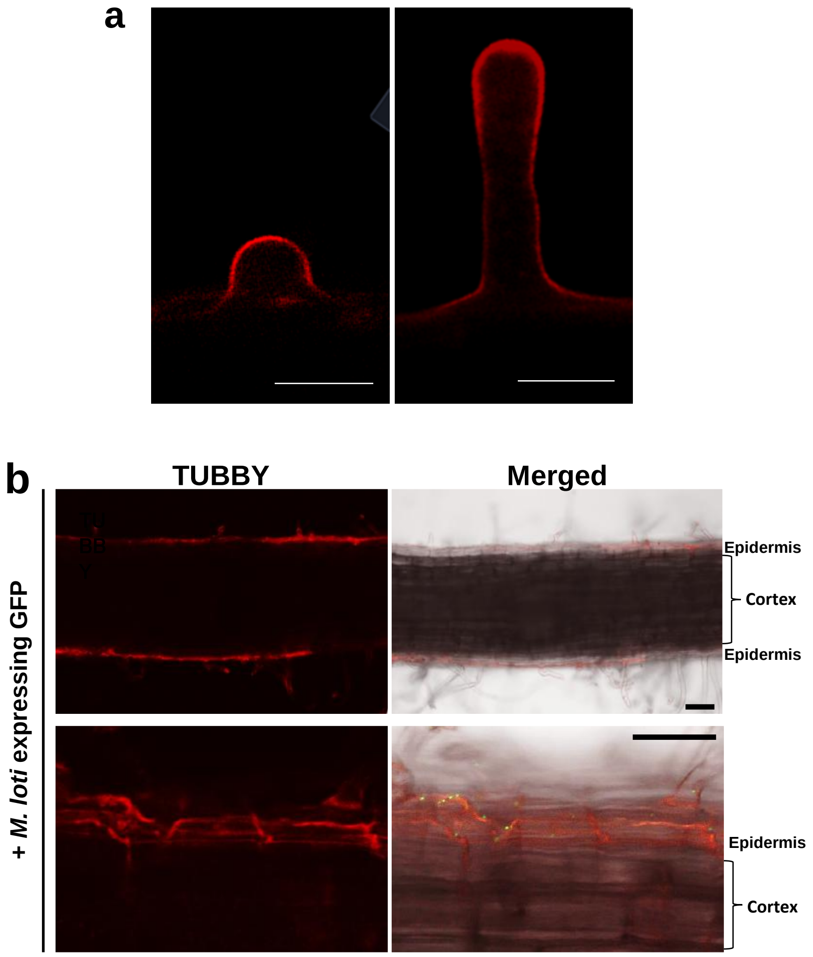


**Supplementary Figure14** *L. japonicus* MG-20 wild-type expressing TUBBY, a PI(4,5)$P_2$ marker, was infected with *M. loti* expressing GFP. Elongating root hair cells (a), the epidermal and cortical cells (b). Scale bar is 20 µm (a), 50 µm (b).

### Supplementary Table 1

| No. of primer sets | Gene or Name | Primer sequences (Fw/Rv) 5' to 3' | Use and specification |
|---|---|---|---|
| 1 | pLjPLP4 Fw 3kb | caccTCATGGATTAGGAAAAATGAAAATG | For cloning of LjPLP4 promoter in pENTR-D-TOPO vector |
| | pLjPLP4 Rv | acatGGTGTCGGTTGGATTGGTTTG | |
| 2 | pLjPIP5K4 Fw 3kb | CGCGGCCGCCCCCTTcaccGAAATAAGAAAAATAGAAAATTG | For cloning of pLjPIP5K4 promoter in pENTR-D-TOPO vector |
| | pLjPIP5K4 Rv | GGCGCGCCCACCCTTTGTGATTTCTTAACTCTTCT | |
| 3 | pLjPIP5K6 Fw 3kb | CGCGGCCGCCCCCTTATCTACAAAACATTTATTG | For cloning of pLjPIP5K6 promoter in pENTR-D-TOPO vector |
| | pLjPIP5K6 Rv | CGGCGCGCCCACCCTTtcatTCTCTTGAAGGTCTCCACTTTG | |
| 4 | LjPLP4 Fw_Venus | GTGACCGCCGCCCTCATGTCTGAACTTCTCTCTTC | For cloning of LjPLP4 gene in pENTR-D-TOPO vector |
| | LjPLP4 Rv | TCGGCGCGCCCACCCTTTTAACAGCAGAAAAACGTCT | |
| 5 | LjPLP4 Fw_mCherry | gacgagctgtacaagATGTCTGAACTTCTCTCTTC | For cloning of LjPLP4 gene in pENTR-D-TOPO vector |
| | LjPLP4 Rv | TCGGCGCGCCCACCCTTTTAACAGCAGAAAAACGTCT | |
| 6 | LjPIP5K6 Fw | CGCGGCCGCCCCCTTcaccATGAGCAGAGAGCTGAATGG | For cloning of LjPIP5K6 gene in pENTR-D-TOPO vector |
| | LjPIP5K6 Rv_Venus | GCCCTTGCTCACCATAGTGTCCTCTACAAAAACTC | |
| 7 | NFR1 Fw | CGCGGCCGCCCCCTTCACCATGAAGCTAAAAACTGGTCTAC | For cloning of NFR1 gene in pENTR-D-TOPO vector |
| | NFR1 Rv_mCherry | CCCTTGCTCACcatTCTCACAGACAGTAAATTTATG | |
| 8 | TOPO Fw | AAGGGTGGGCGCGCCGA | For amplification of pENTR-D-TOPO vector containing Venus |
| | Venus Rv | GAGGGCGGCGGTCACGAACT | |
| 9 | TOPO Fw | AAGGGTGGGCGCGCCGA | For amplification of pENTR-D-TOPO vector containing mCherry |
| | mCherry Rv | GAGAAGTTCAGACATcttgtacagctcgtccatgccg | |
| 10 | Venus_Fw | ATGGTGAGCAAGGGCGAGGAGCTGT | For amplification of pENTR-D-TOPO vector containing Venus |
| | TOPO Rv | AAGGGGGCGGCCGCGGAGCCT | |
| 11 | mCherry_Fw | atggtgagcaagggcgaggag | For amplification of pENTR-D-TOPO vector containing mCherry |
| | TOPO Rv | AAGGGGGCGGCCGCGGAGCCT | |
| 12 | LjPLP4 Fw_pET28 | cagccatatggctagcATGTCTGAACTTCTCTCTTC | For cloning of LjPLP4 gene in pET28 vector |
| | LjPLP4 Rv_pET28 | CGAGTGCGGCCGCGAACAGCAGAAAAACGTCTTCTTC | |
| 13 | pET28_Fw | TCGCGGCCGCACTCGAGAATG | For amplification of pET28 vector |
| | pET28_Rv | gctagccatatggctgccgc | |
| 14 | delta SEC14 Fw | GAGTTTGAATTCAAAGATCAAGGAGGCTGTATGCG | For deletion of SEC14 domain |
| | delta SEC14 Rv | TTTGAATTCAAACTCCTCCA | |
| 15 | qPCR_LjUbi_Fw | ATGCAGATCTTCGTCAAGACCTTG | For qRT-PCR |
| | qPCR_LjUbi_Rv | ACCTCCCCTCAGACGAAG | |
| 16 | qPCR_PpEF1a-Fw | CAAGATCCCGTTCGTGCCTA | For qRT-PCR |
| | qPCR_PpEF1a-RV | GCGTTCAGGTTGTCAAGAGC | |
| 17 | qPCR_CRE-RS1_Fw | TGCAAGTGTCGATGCTCATAGCAATG | For qRT-PCR |
| | qPCR_CRE-RS1_Rv | GATGTTTTGCTGAACCAAGGGATATTG | |
| 18 | qPCR_LjPLP4_Fw | CTCATGGGGAGGAAGAGAAA | For qRT-PCR |
| | qPCR_LjPLP4_Rv | GAGAAACCTGTGGGGTGAGA | |
| 19 | qPCR_LjPIP5K4_Fw | TGGGCAACCTCTTCTGTTCT | For qRT-PCR |
| | qPCR_LjPIP5K4_Rv | TCTCGGTCTCAGGCTTGTCT | |
| 20 | qPCR_PIP5K6_Fw | CATGTTGGTGGGTCTTCACTT | For qRT-PCR |
| | qPCR_PIP5K6_Rv | GGTGCAAGCTCCTGAATAGC | |
| 21 | PLP CRIS1 Fw | attgGAACTTCTCTCTTCAGGGAC | For cloning of the single guide RNA (sgRNA) into pUC19_AtU6oligo vector |
| | PLP CRIS1 Rv | aaacGTCCCTGAAGAGAGAAGTTC | |
| 22 | PLP CRIS2 Fw | attgAGGAATTCTTTGACGAGAAA | For cloning of the single guide RNA (sgRNA) into pUC19_AtU6oligo vector |
| | PLP CRIS2 Rv | aaacTTTCTCGTCAAAGAATTCCT | |
| 23 | PIP5K4 CRIS1 Fw | aatgATACATGGTGTGTCGTCCCG | For cloning of the single guide RNA (sgRNA) into pUC19_AtU6oligo vector |
| | PIP5K4 CRIS1 Rv | aaacCGGGACGACACACCATGTAT | |
| 24 | PIP5K4 CRIS2 Fw | aatgGGAGAATGTCAGTAGATGGG | For cloning of the single guide RNA (sgRNA) into pUC19_AtU6oligo vector |
| | PIP5K4 CRIS2 Rv | aaacCCCATCTACTGACATTCTCC | |
| 25 | PIP5K6 CRIS1 Fw | aatgTATGTTGGTGAATGGAGCAG | For cloning of the single guide RNA (sgRNA) into pUC19_AtU6oligo vector |
| | PIP5K6 CRIS1 Rv | aaacCTGCTCCATTCACCAACATA | |
| 26 | PIP5K6 CRIS2 Fw | aatgGGTTTGGAGTAAAGACCCAA | For cloning of the single guide RNA (sgRNA) into pUC19_AtU6oligo vector |
| | PIP5K6 CRIS2 Rv | aaacTTGGGTCTTTACTCCAAACC | |

### Supplementary Table 1

| No. of primer sets | Gene or Name | Primer sequences (Fw/Rv) 5' to 3' | Use and specification |
|---|---|---|---|
| 27 | U6woISceIwoL1_Fw178 | CTTTTTTTCTTCTTCTTCGTTCATACAG | For amplification of sgRNA cassette |
| | U6woISceI_Rv835 | ttaattaaGGTACCGAGCTCGGATC | |
| 28 | U6woISceIwoL1_Fw178 | CTTTTTTTCTTCTTCTTCGTTCATACAG | For amplification of pUC19_AtU6oligo vector containing sgRNA cassette |
| | U6_Rv77 | ATCGAATTCCCGCGGCCGCTCGA | |
| 29 | PLP4_Cris Fw | GATTGATCATGTTGTTATGGTTGCAGG | For genotyping PCR of plp4 plants |
| | PLP4_Cris Rv | CCTCCACAATAGTGTCAGCAC | |
| 30 | PIP5K4_Cris Fw | TTGGTATGAAGGAGAGTGGA | For genotyping PCR of pip5k4 plants |
| | PIP5K4_Cris Rv | TGAGCCACACTTTCAGAGTTTG | |
| 31 | PIP5K6_CRIS Fw | TCCCCAATGGTGACTTCTAC | For genotyping PCR of pip5k6 plants |
| | PIP5K6_CRIS Rv | CAACTCCACAGAGAACACCTC | |
| 32 | Hyg_Fw | ATGAAAAAGCCTGAACTCACCGCGA | For genotyping PCR of LjPLP4 OE plants |
| | Hyg_Rv | CTATTTCTTTGCCCTCGGACGAGT | |
